# Tissue-wide Erk oscillations set the pace of zebrafish notochord segmentation

**DOI:** 10.1101/2025.08.28.672946

**Authors:** Priyom Adhyapok, James Norman, Michel Bagnat, Stefano Di Talia

**Affiliations:** Department of Cell Biology, Duke University Medical Center, Durham, NC 27710 USA; Center for Quantitative Living Systems, Duke University Medical Center, Durham, NC 27710 USA; Department of Orthopaedic Surgery, Duke University Medical Center, Durham, NC 27710 USA

## Abstract

The generation of a segmented body plan is a hallmark of vertebrate development. Biological oscillators provide a mechanism for timing the formation of repeated structures, yet to date only a few examples of signaling oscillators have been identified in development. Here, we show that the periodic addition of mineralizing segments in the zebrafish notochord is timed by tissue-wide oscillations of Erk activity. We report delayed oscillations in the transcription of *spry* and *dusp* genes, suggesting that they act as negative feedback. Erk activity is regulated by EGFR signaling and increased *egf* expression marks the emergence of oscillations via an infinite-period bifurcation, revealing the mechanism controlling the onset of notochord segmentation and the tunability of the oscillator. Our results show a remarkable degree of synchronization across the notochord and reveal an instance of biological clocks timing a patterning process, crucial for the development of the vertebral column from the notochord.

## Introduction

The generation of periodic structures is a hallmark of embryogenesis; it allows the partition of tissues into units and is a crucial mechanism for sequential patterning of the anterior-posterior axis in vertebrates. Processes like somitogenesis which set up the periodic muscle blueprint early on in development have been shown to engage mechanisms like intra-cellular oscillators whose coordinated actions in space and time help specify segment boundaries^1–4^. However, the formation and positioning of vertebrae is controlled many days later by morphogenetic processes that remain poorly understood^5^. Work in zebrafish has shown that vertebrae assemble using a blueprint generated by the segmentation of the sheath cells surrounding the notochord^6–9^. This process begins around 5 days post fertilization (dpf) and continues for 2 weeks, resulting in the sequential segmentation of the tissue into mineralizing and inter-segment domains while the fish elongates between 4-5 mm. The sequentiality of the process and the repeated addition of new segments requires accurate spatiotemporal control of biochemical activities across multiple tissue scales. While boundaries established by somitogenesis by 1 dpf, and propagation of BMP signaling may explain aspects of sequential segment positioning^10–12^, it remains unclear what sets up the tempo of notochord segmentation allowing it to slowly build this template. Here, we provide evidence that the underlying time-keeping mechanism is driven by oscillations of Erk activity, within the notochord sheath itself. These oscillations show a high similarity across the notochord. We speculate that segmentation relies on themes of synchronization more commonly than previously shown during development.

## Results

### Segmentation of the notochord occurs periodically

To quantitatively capture the dynamic features of notochord segmentation, we performed live imaging in larvae beginning at 7-8 dpf (larvae length: 4-4.25 mm). The analysis was done over a period of around 30 hours, covering around 1mm of the notochord and with images acquired roughly every 4 hours. In the notochord, the expression of the mineralization marker *entpd5a* coincides with activation of Notch signaling^7^. We used a transgenic reporter (*entpd5a:pKRed*) to monitor segment progression (Figures 1A and S1A). To this end, we computationally extracted *entpd5a+* cells (Figure 1A) and tracked fluorescence levels across individual segments over time. We found segments mature by the combined differentiation of sheath cells into *entpd5a+* cells and an increase in fluorescence of previously differentiated cells in segments. Moreover, anterior segments grew more than posterior segments during the period, pointing to the sequential nature of the segmentation pattern (Figures 1A-1C). This also indicates that development of segmented domains occurs gradually along the anterior posterior axis. Given that segmenting regions continue to mature over time, we investigated *entpd5a* dynamics by quantifying the total fluorescence of all detected cells across the notochord (Figure 1D). Monitoring differences in total fluorescence levels between subsequent time points (Figure 1E, segment data in Figure 1F) suggested that increases in *entpd5a* expression appear in periodic bursts, occurring on average every 10-12 hours under our imaging conditions.

**Figure 1.**
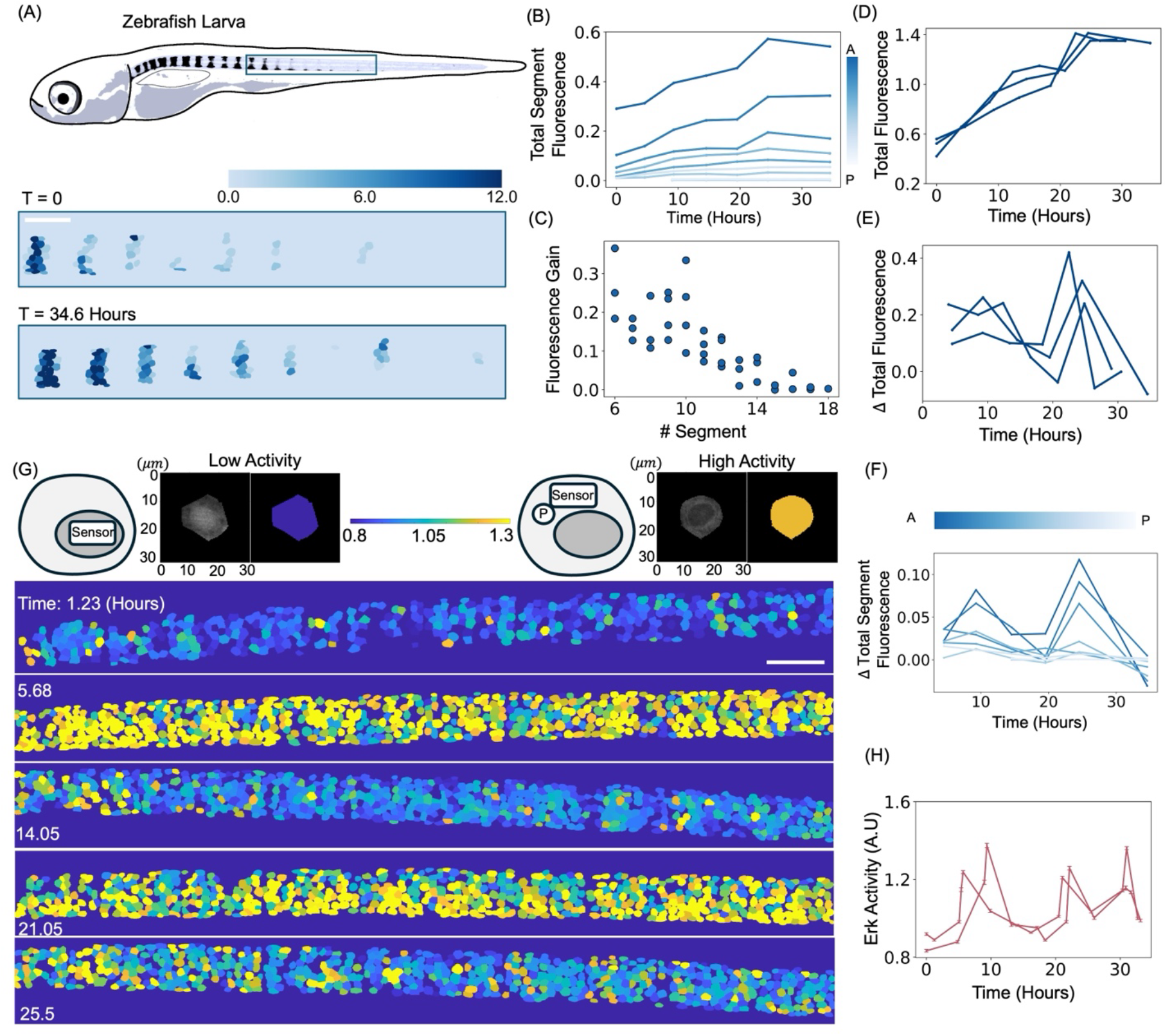
Oscillations of Erk activity in the zebrafish notochord. (A) Top, Schematic of a zebrafish larva (4-4.25mm). Bottom, images showing the initial and final image from a time course over 30 hours demonstrating *entpd5a:pKRed* expressing cells, color coded by their mean fluorescence levels. Scale bar 100 *μm*. (B) Total fluorescence of *entpd5a:pKRed* in segments as a function of time (n = 1, number of segments = 9). Different lines indicate different regions along the AP axis, as shown by the color bar. (C) Fluorescence gain per segment as a function of segment number ordered from Anterior to Posterior, calculated across multiple larvae (n = 4 larvae from N = 3 independent experiments). (D) Total fluorescence levels in the notochord, normalized by mean expression over time per larva show discrete jumps in expression. (E) Timepoint-by-timepoint difference in total fluorescence levels shows bursts in time, indicating periodic activation (n = 3 larvae from N = 3 independent experiments in D-E). (F) Timepoint-by-timepoint differences in *entpd5a:pKRed* fluorescence as a function of time plotted for different regions along the AP axis, as shown by the color bar (n = 1, number of segments = 9). (G) Top, cartoons showing how phosphorylation affects the localization of the sensor along with images and quantification of two individual cells at low and high activity (see colorbar). Bottom, Heat maps displaying Erk activity, reported by *cmn-hsp70l:erk-ktr-eGFP*, quantified in individual cells. Scale bar 100 *μm*. These maps show oscillatory behaviors in the notochord. (H) Average Erk activity ± SEM as a function of time in 2 representative larvae confirms the oscillatory behavior. See also Figure S1.

### Erk activity oscillates across the entire notochord

To identify the molecular mechanism controlling the periodic nature of segmentation, we postulated that segmentation is controlled by a clock driving intracellular oscillations, similarly to what is observed in other systems undergoing periodic patterning^1,13–18^. We suspected that the MAPK/Erk pathway might contribute to the timing of segmentation, as this pathway can generate oscillations in certain developmental and regenerative contexts^19–22^.

To visualize and quantify the activity of the MAPK/Erk pathway in the zebrafish notochord, we generated new transgenic lines driving the expression of an Erk sensor^22–23^ from the *calymmin* locus or from a *collagen 9a2* regulatory sequence, which are specifically expressed in the notochord sheath^24–25^. The Erk sensor is a translocation reporter, in which phosphorylation favors cytoplasmic localization of the reporter. Thus, Erk activity can be estimated by the ratio of cytoplasmic and nuclear fluorescence (schematic in Figure 1G). These transgenic tools allowed us to visualize and quantify the dynamics of Erk activity in the notochord sheath with cellular resolution. Computational segmentation of individual cells over time (Methods, Figures S1B-S1E) revealed that Erk activity is highly dynamic (Figure 1G, Video S1, Figure S1F), with levels in the notochord periodically transitioning between low and high activity. This is evidence of intracellular oscillations in the zebrafish notochord (Figure 1H, n = 2 representative larvae, all data in Figure S1G). We used two metrics to quantify the oscillations: average Erk activity across all the cells (Figure 1H) and percentage of cells that have high Erk activity at a given time (ratio of cytoplasmic to nuclear fluorescence > 1) (Figure S1H). Both metrics revealed clear oscillations and allowed us to estimate a mean time-period for the oscillations of 10 ± 2 hours (Figure S1I). Oscillations are seen in unsegmented and segmenting regions with similar behavior (Figures S2A and S2B). Consistently, we estimated the ratios of time periods in individual segments compared to the rest of the tissue to be 1 (Figure S2C).

Visual inspection of activity heat maps suggests that Erk oscillations extend across the entire notochord sheath, and that the tissue attains similar activity levels across our imaged region. Specifically, we could identify images in which the fraction of cells with high activity, in 45 *μm* bins along a 1 mm region of the Anterior Posterior (AP) axis, ranged from 0.8-1.0 (Figure 2A). Quantifying oscillations in individual bins over time, we found that there were no large-scale tissue differences in the oscillator phase at these sampled times (Figure 2B). To describe the observed synchrony of Erk oscillations, we used the Hilbert transform^26^ to infer the phases of the oscillations across the AP axis of the notochord. We then used these inferred phases to calculate the Kuramoto parameter, a standard mathematical tool to describe the synchronicity of oscillators^27^. For systems oscillating out of synchrony, the Kuramoto order parameter r (Eq 1, Methods) is close to zero, while for synchronous oscillators, the order parameter has a value close to 1. Graphically, this can be visualized by plotting the phases of each binned region on the unit circle and seeing if the phases are clustered (synchronous oscillators) or dispersed across the circle (non-synchronous oscillators). Our data showed strong clustering and allowed us to estimate that the Kuramoto order parameter is typically above 0.9, indicating a high degree of synchrony in the entire notochord (Figures 2C and 2D).

**Figure 2.**
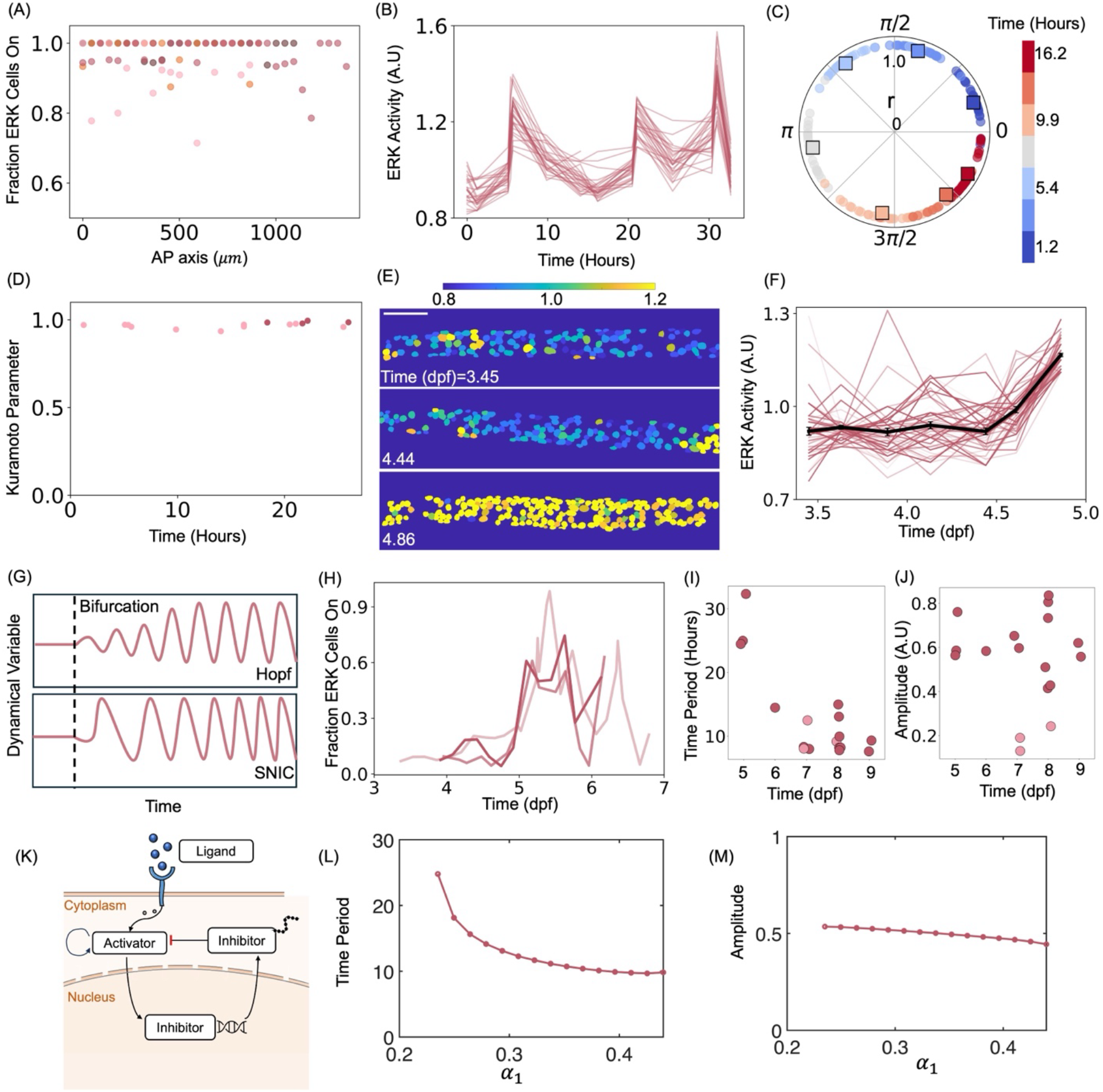
Erk Oscillations are synchronous across the entire notochord and are born via a SNIC-bifurcation. (A) Fraction of cells with high activity quantified in 45 *μm* bins along the AP axis over a 1 mm region of the notochord (colors indicate different larvae, n = 4 larvae from N = 4 independent experiments). This analysis identifies time-points in which a large fraction of cells is active throughout the tissue. (B) Erk activity as a function of time with each line representing a binned region. (C) Estimated phases from each binned regions plotted on the unit circle. Different colors indicate different timepoints in the oscillations (see colorbar). The Kuramoto order parameter (r) for each time point is shown as a square of the appropriate color. The amplitude of the Kuramoto order parameter is given by the distance to the origin, and its phase by the angle along the circle. (D) The amplitude of the Kuramoto order parameter across time (n = 2 larvae from Figure 1H) shows synchronous behavior. Colors indicate the different larvae. (E) Heat map of Erk activity at three different time points between 3-5 dpf. Scale bar 100 *μm*. Transition to high Erk activity seen around 5 dpf (see colorbar). (F) Erk activity (measured in a larva expressing the *col9a2:erk-ktr-mCerulean* reporter) as a function of time in individual cells (n = 1, number of cells = 50, black line indicates average Erk activity ± SEM). Analysis shows the emergence of high Erk activity in the notochord around 5 dpf. (G) Expected dynamic behaviors for different types of bifurcation leading to an oscillatory system. (H) Fraction of cells with high Erk activity as a function of time (n = 3 larvae from N = 3 independent experiments). This analysis shows a bifurcation into oscillatory state, reported by *cmn-hsp70l:erk-ktr-eGFP* and *col9a2:erk-ktr-mCerulean*. (I) Oscillation time-period as function of developmental time. Data were pooled from larvae imaged beginning at different stages of development (Figures S1H and 2H). (J) Amplitude of the oscillation as a function of developmental time for the same larvae shown in (I). Note that the amplitude is in large part similar across all stages, with few outliers showing reduced amplitude, likely due to poor sampling near the oscillation peak given our time resolution. Points colored in pink have less than 70% of cells on at detected peak. (K) Cartoon representing the molecular interactions described by our mathematical model. (L-M) Dependency of the time period and amplitude of oscillations on the bifurcation parameter, *α*_1_ respectively. See also Figure S2.

### Tissue-wide emergence of Erk oscillations

Next, we asked how oscillations emerge in the tissue by investigating the regulation of Erk activity oscillations at times preceding 7 dpf, when 8-9 segments have already begun to form. By imaging larvae every few hours for 2-3 days beginning at 3 dpf, we found that Erk activity is initially low prior to 5 dpf, but transitions to high activity (Figures 2E and 2F, Video S2), in a synchronous manner across individual cells, indicating that the system has undergone a bifurcation. Furthermore, analysis of individual trajectories by monitoring cells crossing a threshold of Erk activity, suggested that Erk dynamics is excitable prior to the bifurcation, as cells getting past the threshold return to low Erk activity only after having reached a significantly higher and transient peak activity level (Figure S2D). To investigate the properties of this transition, we considered two main types of bifurcations typically observed in systems undergoing a transition from a non-oscillatory to an oscillatory pattern,^28–32^ – namely, the super-critical Hopf bifurcation and the Saddle Node on an Invariant Cycle (SNIC, also known as infinite-period) bifurcation. These two bifurcation types are distinguished by how the amplitude and frequency behaves as the system transitions from one state to the other. The Hopf bifurcation is characterized by increasing amplitude and fixed frequency, while the SNIC bifurcation is characterized by fixed amplitude and increasing frequency (Figure 2G). Distinguishing between these two bifurcations provides insight on the properties of the system near transition, specifically on whether the build-up to rhythmicity is gradual, frequency tunable and also informs if the range of response is fixed or variable. Our analysis is consistent with the notochord sheath undergoing a SNIC bifurcation, as the first Erk oscillation is substantially longer (∼24 hours) than subsequent ones (∼10-12 hours) (Figure 2H, 2I and S1I), while the amplitude is similar to the one observed for later oscillations. We note that a few larvae displayed lower amplitude at 7-8 dpf (Figure 2J). However, for these larvae, we could not identify a time point where the fraction of cells with high Erk was larger than 0.7. Since we have always observed such state of high activity in larvae where we had increased our sampling frequency, we speculate that the lower amplitude observed for those larvae at 7-8 dpf is due to limited sampling. Collectively, these observations indicate that the zebrafish notochord displays oscillations that are born via a SNIC bifurcation.

### Delayed oscillations in the expression of *spry* and *dusp* genes

To gain insight on the potential mechanisms controlling the Erk oscillations and the observed bifurcation, we built a mathematical model that can generate oscillations through a SNIC bifurcation (Figures 2K-2M). This model assumes an increase over time in the expression of an Erk activator and the presence of delayed negative feedback, generated by Erk activation of its own inhibitors. Analysis of time-courses with faster sampling near the oscillation peaks identified rapid spikes in Erk activity (Figure S2E). Thus, similar to other models^22^, we included a positive feedback loop to facilitate oscillations and model an oscillatory cycle with multiple time scales as can be seen in systems containing positive and negative feedback loops^33–35^ (Figure S2F).

Given the oscillation period of several hours, we speculated that the negative feedback postulated by the model is provided via transcriptional regulation, and that delays in protein production result in delayed oscillation cycles of the inhibitors relative to Erk (Figures 2K, S2F). To test this hypothesis, we first reanalyzed a transcriptomic dataset^7^ from 13 dpf larvae to identify putative inhibitors expressed in the notochord. In this dataset, notochord sheath cells were FACS-sorted into three separate populations: unsegmented, transitional or segmented domains. We found evidence for the expression in differential spatial patterns of many sprouty (*spry*) and dual-specificity phosphatases (*dusp*) genes, known Erk transcriptional targets and negative regulators of the MAPK pathway^36–38^. To determine how components of this inhibitory network might be regulated in the notochord, we decided to focus on a candidate gene (*spry2*) whose expression is enriched in the unsegmented region (Figure S3A) and would therefore be initially present throughout the notochord. We generated a destabilized transcriptional reporter - *TgKI(spry2-hsp70l:venus-Pest)* - that allowed us to visualize endogenous expression in zebrafish larvae (Figure S3B). We found that *spry2* is expressed in multiple cell types in larvae. Thus, we developed a computational pipeline for the identification of the notochord sheath and for the extraction of any *spry2+* cells in the notochord (Figure S3C). Using this transcriptional reporter and computational analysis, we first tested if *spry2* production is dependent on Erk. For this we used the Mek inhibitor PD0325901, which can effectively reduce Erk activity (Figures S3D-S3F). We treated 8 dpf larvae expressing *spry2-hsp70l:venus-Pest* with either PD0325901 or DMSO for about 24 hours (a period chosen to avoid any confounding effects from potential oscillations and reduced dynamic range). We found that post treatment, *spry2* fluorescence is reduced in the treated larvae as compared to the controls, indicating that *spry2* expression is indeed influenced by Erk activity (Figures S3G and S3H). We then tracked single cells through a time course and found that *spry2* expression oscillates in the notochord sheath (Figures 3A-3C and S3I-S3L). The *spry2* oscillations have similar period to the Erk oscillations and are concurrent with bursts in *entpd5a* expression (Figure 3C). Similar analysis revealed that the peak of Erk activity precedes *entpd5a* expression by few hours supporting the prediction that *spry2* expression is delayed relative to Erk activity (Figures S9B and S9C; the maturation time and lifetime of Venus-Pest likely contribute to part of the estimated delay). Consistent with the transcriptomic results, we find that as segmentation proceeds, the expression of the *spry2* reporter is downregulated in the segmenting regions (Figures S4A and S4B), as shown by extracting the average expression levels in segmenting and non-segmenting cells (Figure S4C). To test if other Erk inhibitors are correspondingly upregulated in the mineralizing regions, we examined *dusp6,* which our transcriptomic analysis revealed to be enriched in the *entpd5a*+ domains (Figure S4D). To monitor *dusp6* expression, we generated a similar transcriptional reporter - *TgKI(dusp6-hsp70l:venus-Pest)* (Figure S4E). Analysis of the spatial patterns confirmed that *dusp6* expression closely aligns with *entpd5a*+ cells (Figures S4F-S4H). To verify that *dusp6* expression is similarly dependent on Erk, we blocked activity in 8 dpf larvae with a Mek inhibition experiment for about 24 hours. After extracting any detectable individual cells in the PD0325901 treated larvae, we found reduced levels of expression as compared to the DMSO controls (Figures S5A and S5B). Next, after tracking individual segmenting cells for about 30 hours, we found that expression of *dusp6* increased over time. This increase was marked by periodic bursts (Figures 3D and 3E). Facilitated by the timing of these bursts, we estimated local baselines to extract the underlying oscillatory patterns in this reporter (Figures S5C-S5J). These changes in *dusp6* expression demonstrate a cycle duration and peak times similar to those of bursts in *entpd5a+* addition (Figure 3F), suggesting that, similarly to *spry2*, *dusp6* expression might be delayed relative to Erk. Together, our observations on two candidate genes support a model in which the *spry* and *dusp* inhibitory networks are expressed in an oscillatory manner, delayed relative to Erk. Additionally, simultaneous knock down of both *spry2* and *dusp6* showed a reduced rate of segment progression (Figures S6A-S6D) and reduced range of Erk activity (Figures S6E-S6H) showing less effective inhibition of Erk. These observations suggest that these factors contribute to inhibition of Erk activity, likely in combination with other *spry* and *dusp* genes.

**Figure 3.**
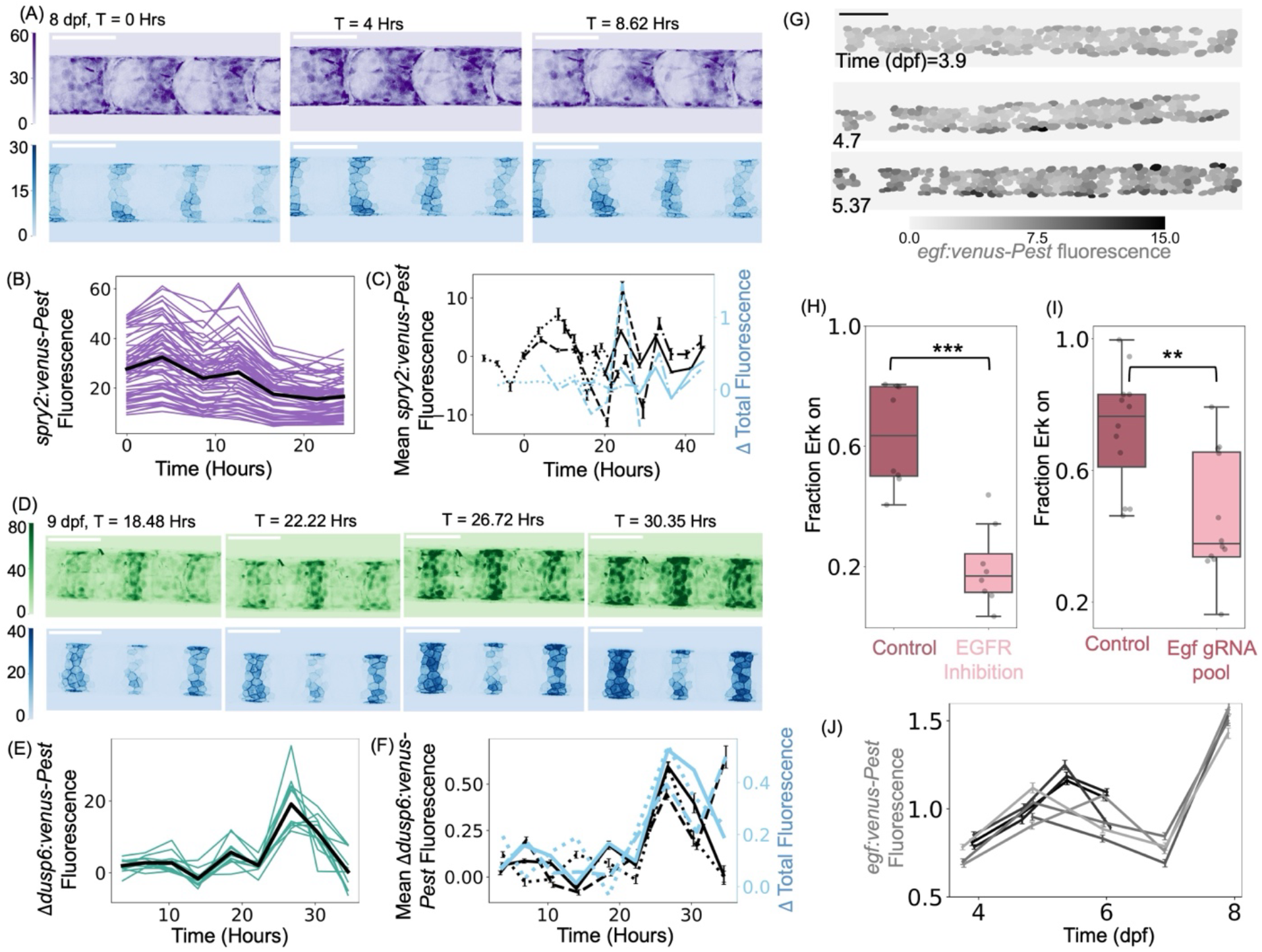
Identification of players involved in the MAPK/Erk pathway oscillator. (A) Heatmaps at 8 dpf visualizing *spry2-hsp70l:venus-Pest* and *entpd5a:pKRed* fluorescence over time with intensity values mapped by the respective colorbars. Scale bar 100 *μm*. (B) *spry2-hsp70l:venus-Pest* fluorescence as a function of time for single cell tracks from an 8 dpf larva (n = 1, number of cells = 69). This analysis demonstrates oscillations. (C) Detrended *spry2-hsp70l:venus-Pest* expression levels along with timepoint-by-timepoint difference in *entpd5a* fluorescence as a function of time. Data is plotted as Mean±SEM and larvae are aligned for visualization (n = 4 larvae over N = 4 independent experiments). (D) Heatmaps at 9 dpf visualizing *dusp6-hsp70l:venus-Pest* and *entpd5a:pKRed* fluorescence over time with intensity values mapped by the respective colorbars. Scale bar 100 *μm*. (E) Timepoint-by-timepoint differences in *dusp6* fluorescence expression (n = 1, number of cells = 12) as function of time in an 8 dpf larva. Black line indicates the average over all the traces. (F) Mean timepoint-by-timepoint differences±SEM in *dusp6* expression along with differences in *entpd5a* fluorescence as a function of time (n = 3 larvae shown from one experiment). (G) Segmented cells detected by *egf-hsp70l:venus-Pest* fluorescence in a single larva over time, color coded by mean fluorescence levels per cell. Scale bar 100 *μm*. (H) Fraction of cells with high Erk activity at 5 dpf following an EGFR inhibition treatment (N = 2 independent experiments; n = 8 larvae per condition; Mann-Whitney U test; *p* = 0.00031). (I) Fraction of cells with high Erk activity at 5 dpf following injections with an sgRNA pool targeting *egf* (N = 2 independent experiments; n = 12 larvae per condition; Mann-Whitney U test; *p* = 0.0017). (J) Average±SEM *egf-hsp70l:venus-Pest* fluorescence as a function of developmental time. Each line indicates a different larva (n = 7 from N = 2 independent experiments). This analysis indicates an increase in *egf* expression between 4-8 dpf with a drop around 7 dpf. See also Figures S3-S8.

### Expression of *egf* coincides with the onset of Erk oscillations

Next, we tested the model prediction that the SNIC bifurcation requires the increasing expression of an Erk activator across the notochord. Analysis of the transcriptomics data revealed the possible involvement of the Epidermal Growth Factor (EGF) pathway (Figure S7A). Thus, we generated a transcriptional reporter to study the expression of *egf* - *TgKI(egf-hsp70l:venus-Pest)* (Figure S7B). Remarkably, we found *egf* expression increasing in the notochord prior to the SNIC bifurcation at 5 dpf (Figures 3G and S7C), around the time when the first jumps in *entpd5a:pKRed* expression occur, marking the beginning of segmentation (Figure S7D). Increased *egf* expression was independently confirmed by Hybridization Chain Reaction (HCR) in-situ (Figure S7E). Pharmacological inhibition of EGF receptors (EGFR) between 3-5 dpf resulted in a significant decrease of Erk activity across the notochord, at a time when we would have expected Erk activity to be high post bifurcation (Figures 3H and S7F). Similarly, we observed a reduction in the number of Erk active cells in the 5 dpf notochord by CRISPR-Cas-mediated knockdown of *egf* (Figures 3I, S7G). These observations are consistent with our model postulating that increasing expression of Egf ligand can drive a bifurcation in Erk dynamics. We also observe that at 5 dpf, while there is some variability in the cell-to-cell level of expression, *egf* expressing cells are distributed through the entire tissue (Figures 3G, S7C). Thus, an increase in *egf* expression seems to be a major driver of the bifurcation that initiates Erk oscillations.

Next, we investigated *egf* expression following the bifurcation and found that it becomes more complex in both space and time. We detected a transient decrease in *egf* expression in notochord sheath cells at 7dpf, followed by a significant increase again at 8 dpf. We speculate that the drop in average *egf* expression seen around 7 dpf could arise from coupling between the oscillatory network and *egf* expression (Figure 3J), which might also become oscillatory. Moreover, the lifetime of the Egf protein might be such that a transient reduction in its transcription does not trigger the transition back to a non-oscillatory state. By 8 dpf, clear spatial patterns of *egf* expression also emerge (Figures S8A and S8B). Sampling larvae with a higher temporal resolution (time frame ∼30 minutes) close to the time of peak Erk activity revealed spatial patterns of Erk activity with similar features (Figures S8C-S8F). We speculate that the emerging spatial pattern of *egf* expression might set these spatial variations in Erk activity dynamics.

### The timing of Erk activation correlates with the timing of notochord sheath cell differentiation

Having characterized the Erk oscillator, we set out to test its functional significance. The period of the Erk oscillations was very similar to the duration between increases in *entpd5a* expression, suggesting a direct link between Erk oscillations and the development of mineralizing domains in the notochord. Analysis of *entpd5a* expression and Erk activity in the same larvae indicated that the bursts of *entpd5a* expression closely follow Erk activity peaks (Figures 4A-4B and S9A). We estimate a delay of a few hours between Erk activation and the next burst of *entpd5a* expression (Figures S9B and S9C); the maturation time of the reporter likely contribute to the estimated delay. Additionally, we tested whether the onset of Erk oscillations coincides with the onset of sheath segmentation. To this end, we combined both Erk and *entpd5a* dynamics between 3-6 dpf. We observed a rapid increase in the rate of *entpd5a* addition that coincided with the onset of Erk oscillations (Figures 4C and S9D), suggesting that Erk activity controls the timing of segment specification. As such, blocking Erk activity with a pharmacological inhibitor should interfere with segmentation. To test this prediction, we again used the Mek inhibitor PD0325901. Starting at both 5 and 7 dpf, larvae were exposed over 50 hours to 2 *μM* of PD0325901 (Figure 4D). We found that the number of *entpd5a+* cells added in forming segments was significantly reduced (Figure 4E). We also observed a significant reduction in the gain of overall *entpd5a+* fluorescence (Figure 4F), suggesting that in addition to the specification of new *entpd5a+* cells, Erk activity also promotes the continued expression of *entpd5a* in already differentiated cells. Similar results were obtained by blocking EGF receptor signaling at 7 dpf pharmacologically (Figures S9E-S9F). Taken together, these observations support the view that Erk activity is required for segmentation.

**Figure 4.**
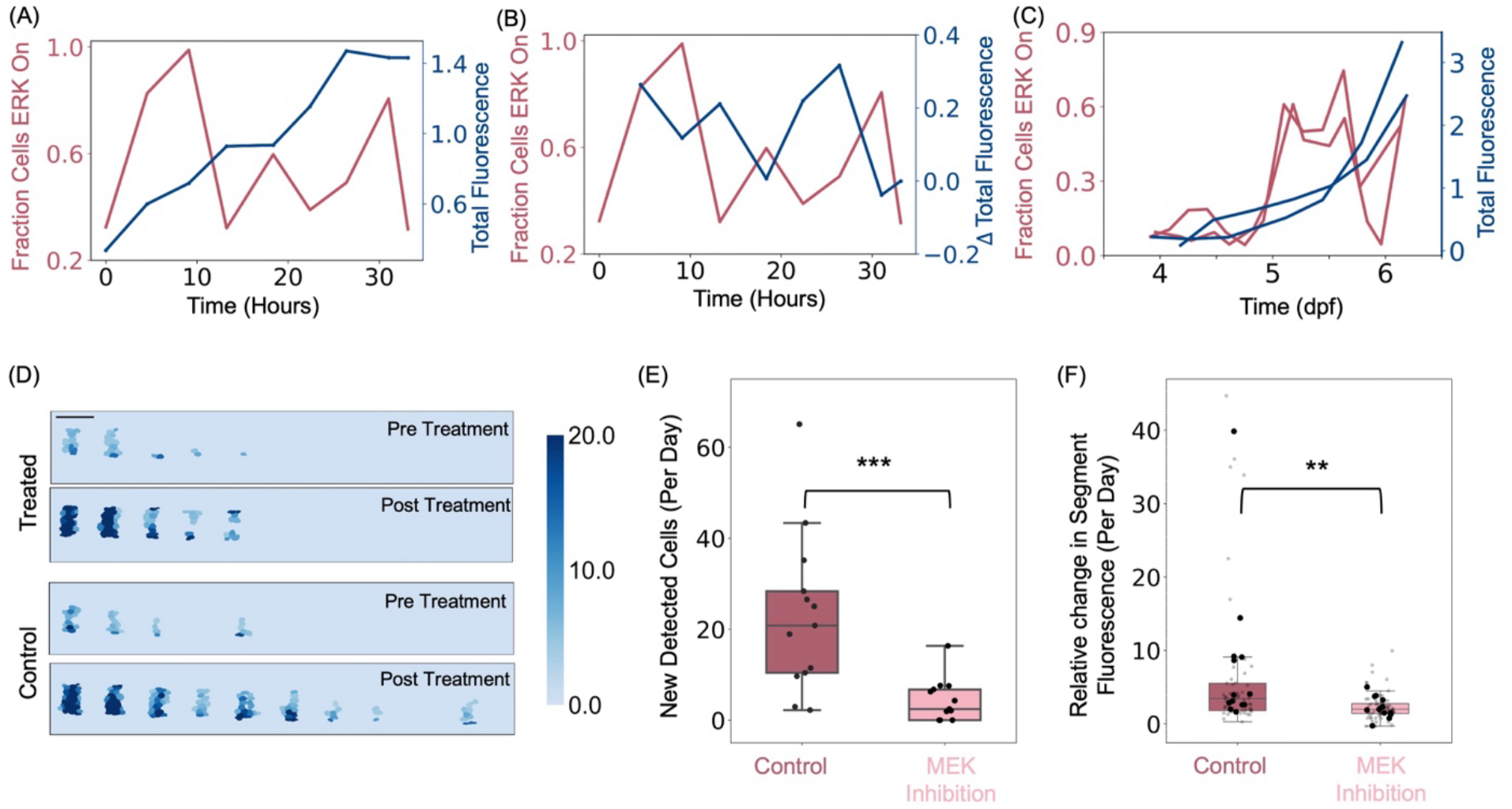
Erk oscillations control the pace of segmentation. (A) Fraction of cells with high Erk activity and total *entpd5a:pKRed* fluorescence normalized by mean levels as a function of time in the same larva. (B) Fraction of cells with high Erk activity and timepoint-by-timepoint difference in *entpd5a:pKRed* fluorescence as a function of time in a single larva. *entpd5a:pKRed* fluorescence bursts are delayed with respect to the oscillation cycle. (C) Fraction of cells with high Erk activity and timepoint-by-timepoint difference in *entpd5a:pKRed* fluorescence levels as a function of developmental time (n = 2 larvae from N = 2 independent experiments). This analysis shows how the first oscillation cycle correlates with the first burst in *entpd5a* expression. (D) Heat maps of *entpd5a* expression quantified in single cells in larvae treated with DMSO or 2 μM Mek inhibitor for 50 hours. Intensity values are mapped by the colorbar. Scale bar 100 μm. (E) Number of newly added cells in control larvae and larvae in which Mek was inhibited pharmacologically; Mann-Whitney U test, *p* = 0.00048. (F) Relative change in segment fluorescence in cells already differentiated in control larvae and larvae in which Mek was inhibited pharmacologically. Small points indicate segment-level measurements, large points indicate per-larva means on which statistical comparison was performed (Mann-Whitney U test, *p =* 0.0077). N = 4 independent experiments with n = 13 larvae per condition for (E-F). Data indicate that Mek inhibition results in fewer newly added cells and less maturation of existing segments. See also Figure S9.

To test whether Erk activation indeed affects differentiation timing, we designed an experiment to block Erk activity transiently and determine whether that induces a predicted delay in the segmentation process. To this end, we selected larvae that on average had high Erk activity and treated them with the Mek inhibitor for about two hours, after which the drug was washed out (Figure 5A). In the control larvae, we were able to identify an Erk peak within the next 10 hours (Video S3); in contrast, Erk activity in treated larvae showed a delay, peaking at around 14 hours (Video S4, Figure 5A and 5B). Similarly, after the treatment, we detected a first burst in *entpd5a*:*pKRed* expression in both experimental groups, albeit with a delay in the treated condition that we attribute to effects of the washout experiment. Consequently, we were able to detect the next burst in *entpd5a*:*pKRed* expression after the peaks of Erk activity in both groups, effectively elongating the duration between the *entpd5a* peaks in the treated larvae as compared to the controls (about 12 hours as compared to 8 hours, similar to the duration of time between the peaks of Erk activity) (Figures 5C and 5D). We additionally performed an experiment to change the number of oscillations (average oscillation frequency) by treating larvae with two 10-hour pulses of Mek inhibition and allowing the system to recover in between over a period of 2 days (Figure 5E). Consequently, after beginning the experiment with similar *entpd5a*:*pKRed* fluorescence levels for both the treated and DMSO control groups (Figure S9G), we detected a reduced pace of mineralization for the treated group (Figure 5F). Collectively, these experiments implicate Erk oscillations in controlling the timing of segmentation of the notochord sheath.

**Figure 5.**
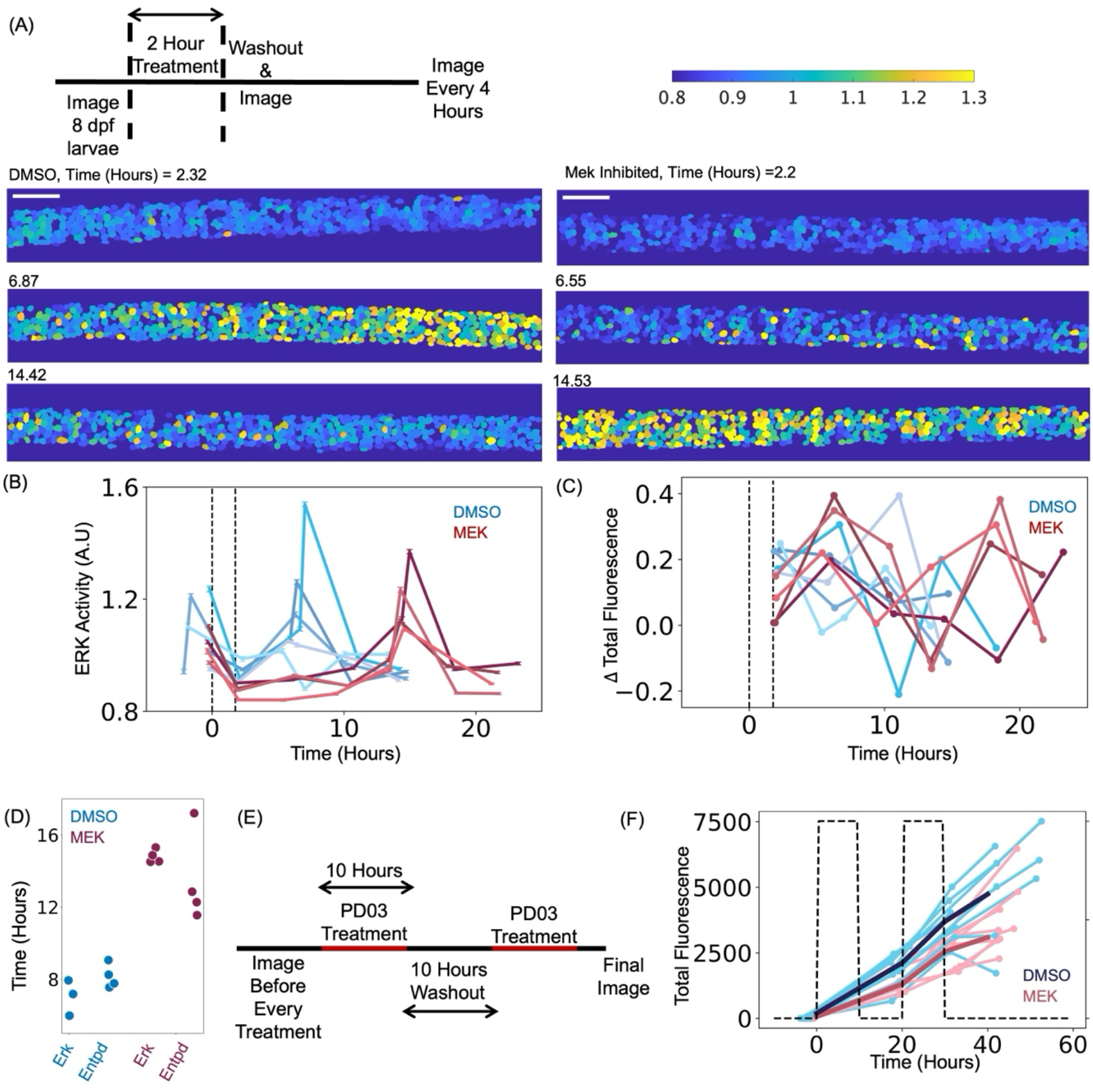
Altering Erk dynamics changes the dynamics of segmentation. (A) Top, Experimental design to alter segmentation dynamics. Starting with similar high Erk activity, 8 dpf larvae were exposed to DMSO or Mek inhibitor for about 2 hours and monitored for an additional oscillation period. Bottom panels, heat maps of Erk activity in DMSO (left) or Mek inhibited (right) at different times following post treatment. Scale bar 100 μm. (B) Erk activity as function of time, dotted lines indicate treatment window. Treated larvae reach a new peak of Erk activity with a delay relative to control larvae (n = 5 controls, n = 4 treated over N = 5 independent experiments). (C) Timepoint-by-timepoint differences in *entpd5a:pKRed* fluorescence levels as a function of time in the same larvae from (B) show bursts in expression following peaks of Erk activity. (D) Time elapsed between the first peak of Erk at the onset of experiment to the detection of next Erk peak following treatment along with time difference between detected bursts of *entpd5a* expression (n = 4 calculated per condition). (E) Experimental design to reduce the number of oscillation cycles through two 10-hour pulses of Erk inhibition. (F) Change in total *entpd5a* over the course of experiment in controls (cyan lines) versus treated (red lines); N = 3 independent experiments, n = 11 control larvae, n = 9 treated larvae. Darker lines indicate medians of the two groups measured after interpolating at intermediate intervals. Dashed black lines indicate the 10-hour drug intervals with the washout period in between. See also Figure S9.

## Discussion

Periodic patterning of the body plan is a hallmark of vertebrates which typically initiates with somite segmentation in early embryogenesis. Following this early developmental process, other mechanisms must operate to ensure that derived structures, such as vertebrae, are properly positioned. The zebrafish notochord provides a great model system to investigate these mechanisms as it was shown that the position of the vertebrae is ultimately specified by a segmentation process that partitions the notochord in alternating segments. This segmentation process starts at 5 days post-fertilization (4 days after somitogenesis is completed) and is influenced by the previous positioning of somites, which ensures the proper positioning of mineralizing segments and thus plays a role in integrating the two patterning processes. Previous experiments had implicated Notch as essential for notochord segmentation^7^. However, the maturation and growth of notochord segments over time suggest that Notch does not play the same role in synchronizing boundary formation as in somitogenesis^39^. Moreover, restriction of Notch activity to segmenting domains and its sustained activation over time (Figures 1A and 1B), which contrasts with the oscillatory expression typical of somitogenesis^13,39^, suggests there are other pathways that are responsible for periodically transforming unsegmented regions into a segment. Thus, we decided to test whether other pathways might provide the oscillatory activity, needed for a clock mechanism timing notochord segmentation. Using an activity biosensor, we have identified oscillations of Erk activity in the zebrafish notochord sheath. The oscillations strongly correlate with the timing of segmentation of the notochord, arguing that they represent the mechanism controlling the tempo at which mineralizing segments are established to build the vertebral column. This is further supported by experiments altering the properties of the Erk oscillations. However, these experiments do not distinguish whether Erk oscillations are required for segmentation or whether the duration of the periods during which Erk is active set the rate of mineralization. Distinguishing between these two scenarios will require in the future the development of more surgical methods for manipulating Erk and EGFR activity. The Erk oscillations are present throughout the tissue, even as segmenting domains are specified. This suggests sequential segment formation may arise from a wavefront of BMP^11^ signaling propagating through the tissue and receiving inputs from the clock controlled by Erk oscillations to activate Notch and the mineralization program at the positions where somite boundaries contact the notochord. As such, notochord patterning resembles the original clock and wavefront model theoretically proposed by Cooke and Zeeman fifty years ago^3^. Thus, dissecting the signaling interactions that ensure sequential, periodic patterning of the notochord provide a unique system to test a new quantitative model of tissue patterning.

Signaling oscillations are ubiquitous in biological systems and provide a huge variety of functions^4^. In these systems understanding the mechanisms by which a biological system can transition from an oscillatory to a non-oscillatory behavior is of crucial importance. Dynamical systems theory provides a clear conceptual guide to understand those transitions by identifying the underlying bifurcations^40^. Here, we have shown that the onset of oscillations occurs through an infinite-period (SNIC) bifurcation^28^, implying that the build-up to rhythmicity is gradual, frequency tunable and the amplitude of response fixed^32^. Through a combination of mathematical modeling and experimental analysis, we argue that the onset of the oscillations is controlled by the increasing expression of the Egf ligand and delayed negative feedback mediated by *spry* and *dusp*. However, this model remains to be fully tested. While we have focused on two candidate genes (*spry2* and *dusp6*), several other *spry* and *dusp* genes are expressed in the notochord and likely to contribute to and/or be required for the Erk oscillations. Similarly, our experiments suggest a more complex spatiotemporal regulation of the expression of Egf ligands and that such regulation might be responsible for spatiotemporal features of Erk oscillations. We recognize that future work will be required to identify all the ligands, receptors and inhibitors controlling the Erk oscillator.

The bifurcation and synchronization^26,41–42^ observed here are features that can explain the initiation and maintenance of notochord segments. We propose that similar features might be important in other systems and, specifically, that the mechanisms that we are identifying in the notochord will provide a conceptual framework for the study of other biological tubes such as neural tube, trachea, and nephrons which show segmented gene expression patterns whose dynamics are not understood^43–44^. Finally, we speculate that the use of biosensors and quantitative methods, similar to the ones we have developed here, will yield more examples of signaling oscillations and synchrony controlling developmental processes.

## Resource Availability

Further information and requests for reagents and resources should be directed to and will be fulfilled by the corresponding authors Michel Bagnat and Stefano Di Talia.

## Data and materials availability

Raw data generated for this study will be deposited publicly. All materials, reagents are available upon request without any restrictions. Main custom codes used in the study have been deposited at https://github.com/padhyapok/notochord_oscillator

## Acknowledgments

We would like to thank Dr. S. Wopat and J. Bagwell for assistance in generating reporter lines. We thank Dr. B. Cox for sharing transgenic fish and T. Hao for help with data analysis. We thank the Duke ZeCore staff and Brennen Kosmeh for zebrafish care. We are grateful to Dr. R. Diegmiller, Dr. B. Hogan, Z. Lu, Dr. B. Mathey-Prevot and Dr. K. Poss for critical reading of the manuscript. This work was funded by the National Institutes of Health grants R01GM150666 (MB) and R01HD119820 (SDT/MB) along with a Duke Science and Technology Spark Award (SDT) and a Stem Cell Award from Shipley Foundation (SDT).

## Author Contributions

Conceptualization: PA, MB, SDT; Data curation: PA; Formal analysis: PA; Funding acquisition: MB, SDT; Investigation: PA, JN; Methodology: PA, JN, MB, SDT; Project administration: MB, SDT; Resources: MB, SDT; Software: PA; Supervision: MB, SDT Validation: PA, JN; Visualization: PA, MB, SDT; Writing – original draft: PA, MB, SDT; Writing – review & editing: PA, MB, SDT

## Methods

### Experimental Model and Study Details

Zebrafish of Ekkwill and AB strains were housed and bred under standard conditions^45^. No statistical methods were used to pre-determine sample size for experiments. Larvae were picked randomly from a clutch at different age stages and assessed for Standard Length^46^, measured in a straight line between the snout to the tip of the notochord. Any larvae with measurements that deviated from the rest of the experimental group were not followed for imaging or subsequent computational analysis. Where possible, additional stage matching was done with the presence of genetic markers that indicated the number of segments. Replicates were performed with independent clutches. Unless otherwise noted, throughout the text N refers to the number of independent experiments (replicates performed with different clutches) and n refers to the number of individual larvae examined within the experiments. Where applicable, equivalent number of larvae were used per replicate, per treatment condition. Researchers were not blinded during data visualization or analysis. All fish experiments were approved by the Institutional Animal Care and Use Committee at Duke University and followed all the relevant guidelines and regulations.

### Generation of transgenic Lines

Our strategy for generating endogenous reporters was adapted from previous studies^11,47^. A donor sequence containing the heat shock promoter (*hsp70l*) and a destabilized form of *Venus* (*venus-Pest*) was generated in a *puc19* plasmid. Additionally, donor sequence containing the heat shock promoter (*hsp70l*) and *erk-ktr-eGFP* was generated in a *puc19* plasmid using In-Fusion cloning (Takara).

Then PCR donor were amplified from the plasmids using the following primers: puc19_F: 5′ GCGATTAAGTTGGGTAACGC 3′; puc19_R: 5′ TCCGGCTCGTATGTTGTGTG 3′.

The following gRNA target sites were used to generate the endogenous reporters: *spry2* (GGAAGGCTGGTCATCAATGGGG), *dusp6* (GCGATCGTCGCCTGGAAACGGGG), *egf* (AGCCTCAAAGTCCTACACCAGGG), *cmn* (TGACACATGCTGTTCCTTATTGG).

A stock injection solution containing the following was injected at the single cell stage: PCR donor (10 ng/μL), Cas9 protein (500 ng/μL), target site gRNA (50 ng/μL), phenol red.

The *erk-ktr-mCerulean* cassette was subcloned from *pSKS-I:osx-erk-ktr-mCerulean*^22^ using the following primers:

F-5’ GGGGACAAGTTTGTACAAAAAAGCAGGCTCCATGAAGGGCCGAAAGCCTCG 3’

R-5’ GGGGACCACTTTGTACAAGAAAGCTGGGTCGATCTAGAGGATCATAATCA 3’

A BP clonase reaction generated a *pME-erk-ktr-mCerulean* plasmid. A subsequent Gateway LR reaction produced the final *pTol2CG2:col9a2-erk-ktr-mCerulean* plasmid.

One-cell stage embryos were co-injected with the Tol2 construct and transposase cRNA.

### Genome editing

Single-guide RNAs (sgRNAs) targeting *spry2* and *dusp6* were designed to introduce insertion/deletion (indel) mutations within coding exons near predicted phosphorylation sites (*spry2* exon1 Y52^36^ and *dusp6* exon2 Ser159^48^) and near the kinase interacting motif (*dusp6* exon 1 Arg64/65^49^). The pooled sgRNA mixture contained the following target sequences:

*spry2:* GGGGGCCGTGGCGCCACCGT

*dusp6:* GGCCCAGGACCTGGGAGGTT; GGTTGCCTTTCTTGAGTCGC

Zebrafish embryos were microinjected at the one-cell stage with an injection mix consisting of Cas9 mRNA (500 ng/μL) and pooled sgRNAs (50 ng/μL each).

Genome editing efficiency was assessed using the heteroduplex mobility assay (HMA). Genomic DNA was extracted from injected and non-injected embryos, and the target loci were amplified using the following primer pairs:

*spry2*

Forward: AGCTTTGCGTGACAGTGGCA

Reverse: GAGCCCTCAGGGAGTTCCTA

*dusp6*

Forward: ATGGACTGCCGAGCGCAAGA

Reverse: GACTCGATGTCCGAGGAGTC

For *egf* knockdown, a pooled sgRNA mixture comprising the following guide sequences was used: gRNA 1 GGCAGGGCCTGAAGCTCGTA, gRNA 2 GGCAGTAACAGGTAACCCTC, gRNA 3 GGCAGTGGTCAGCACCGGTC.

To assess the efficiency of *egf* knockdown, total RNA was isolated from 30 injected and non-injected embryos and reverse-transcribed to generate cDNA using SuperScript III (Invitrogen: 18080-093) following the manufacturers protocol. Expression of *egf* was then evaluated by RT-PCR using the following primers:

Forward: GGACAGATATGGGTGATCGTC

Reverse: GCGATATCAAATGGACGATC

### Microscopy

All microscopy was performed on a Leica SP8 inverted confocal scope equipped with Leica Application Suite. Larvae were anesthetized with 0.04% Tricaine (VWR:MSPP-MS222) and mounted on glass bottom dishes using 0.9% low melt agarose in egg water. Imaging parameters were kept constant across each experiment. Time courses with EGFP, Venus or PKRed fluorescent proteins were performed using a Fluotar VISIR 25x/0.95 water based objective. Experiments with mCerulean were performed with a HC PL APO CS2 40x/1.25 Glycerol objective or HC PL APO CS2 20x/0.75 Immersion Oil based objective. Whole fish images were taken on a HC PL APO CS2 10x/0.40 dry objective. The fluorescent proteins were excited with the following lasers: 405 nm for mCerulean, 488 nm for EGFP/Venus, 552 nm for PKRed. The notochord was divided into individual tiled regions and the region that could be covered, optimized by imaging duration was dependent on the fluorophore. Typical regions of the notochord with transgenic lines containing EGFP, Venus, PKRed fluorophores were 1000 *μm*, while experiments with mCerulean covered around 500 *μm*. This region was divided into individual tiled regions, (e.g a tile using a 25x objective at 3x optical zoom covered 155 *μm*) and then stitched using Leica software unless specified otherwise. Larvae were revived in between time points.

### Data Analysis

#### entpd5a fluorescence extraction

Image segmentation was performed on 2D max projections spanning the Anterior-Posterior (X) axis and the Dorsal Ventral (Y) axis. The surface of the notochord maps out a cylindrical tube with *entpd5a+* cells arranged around in a ring. To avoid overlapping signal in the max projections, the notochord was cut in half along the medial-lateral (Z) axis. This was possible in larvae combined with the transgenic line *TgKI(cmn-hsp70l:erk-ktr-eGFP)*, as the specificity of the fluorescence expression pattern enabled computational extraction of the notochord region. First, for each X position on the AP axis (Figure S1B), the fluorescent EGFP signal in the Y-Z plane was thresholded. Next, a binary mask was generated by removing any pixels beyond the center of the thresholded signal along the Z axis. After multiplying the modified mask with the original fluorescence reported in the Y-Z plane (either for PKRed or EGFP) (Figure S1C), the slices were combined to reconstruct the notochord along the AP axis and finally max projected along the Z axis. Cells were segmented using Cellpose^50^. Total fluorescence from individual cells was used to map expression over time. Special attention was paid at each time while adding new cells to the detected count, especially at the periphery of the DV axis, as remounting the larvae during subsequent time-points might have shifted the orientation of the notochord. Fluorescence intensities were rescaled by the time average for better comparison between fish samples.

Total intensities per segment were calculated by automatically assigning cells to different segment clusters. All individual cell positions along the Anterior Posterior axis were first sorted. Differences in positions with any large breaks helped separate the cells into the clusters. Individual intensities were summed up for the whole segment and the averaged centroid position of the cells was taken to be the segment location. Segments were then tracked by finding the nearest cluster match between time points. Any segment clusters which could not be assigned to a nearest position in the previous image was assigned as a new segment formed.

#### Extraction of Erk activity

Individual EGFP-expressing cells were segmented using Cellpose using models trained on notochord specific datasets in combination with pre-trained models from the Cellpose repository. After manual curation of any incorrect masks, another model was trained to detect nuclei within each cell corresponding to regions of low or high intensity. The efficiency of the model was also tested on a dataset generated manually by calculating intersection over union of the masks (Figure S1D).

The cytoplasmic intensity from the Erk sensor was estimated in a ring, 3 pixels out from the detected nucleus (Figure S1E). This was done to best avoid any overlapping cytoplasmic signal that could arise from neighboring cells after the max projection. Sensor activity was calculated as a ratio of the intensity in the cytoplasm compared to the nuclear intensity.

Erk activity was then averaged across all detected cells over time. We also calculated the fraction of cells whose cytoplasmic levels were greater compared to intensity in the nucleus, indicating a cell with high activity. Time period of oscillations was calculated by measuring trough-trough distances from the time series.

Figures S2A-S2C were generated by identifying larvae which were sampled close to an oscillation peak. For detected PKRed+ cells, Euclidean distances were then calculated to find the closest EGFP+ cell and calculate the corresponding Erk activity. Cells were then assigned into segments and the average Erk activity calculated for the entire segment (Methods: *entpd5a* fluorescence extraction).

For Figure 2I, data from Figure S1H was separated by developmental time and combined with data from Figure 2H.

For larvae expressing mCerulean as a fluorophore, microscopy parameters were optimized to maximize the signal-to-noise ratio over imaging durations that did not impact fish health. Different methods were used to extract cells and nuclei. Cells were extracted either by training a separate Cellpose model or manually. Nuclei were then estimated by training either a separate model or eroding cells by 10 pixels, to approximate the nuclear region. The same analysis was used across a time-course or control vs treated conditions. The data in Figure 2F was collected with larvae crossed to *ubi:h2a-mCherry*^51^. This allowed easier estimation of the nuclear region, but we were limited by the number of nuclei that could be separated out from the surrounding tissues, given the ubiquitous expression of *ubi:h2a-mCherry*.

#### Kuramoto Parameter

Individual images over a time series were first cropped using ImageJ/Fiji to cover the same region. Erk activity was then binned in 45 *μm* bins along the AP axis (containing on average of 18 cells). After identifying an oscillation cycle using the average activity for the whole notochord, a Hilbert transform was applied to each binned trace (*j*) to extract a phase *θ*_j_. The phases were combined over *N* bins using the equation:

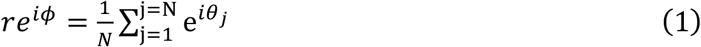

Finally, the average phase (*φ*) and amplitude (r) were plotted on the unit circle along with the phase distributions at each time point.

#### Mathematical Modeling

Our model is composed of ODEs representing activator-inhibitor interactions. We model a negative feedback loop with inhibitor suppressing activator production and a positive feedback loop for the activator, modeling a term where the activator enhances its own production. Combination of positive and negative feedback loops have been used to study other oscillatory systems^22,52^ and methods for modeling oscillators have been reviewed in other texts^53^.

Our equations have the following form -

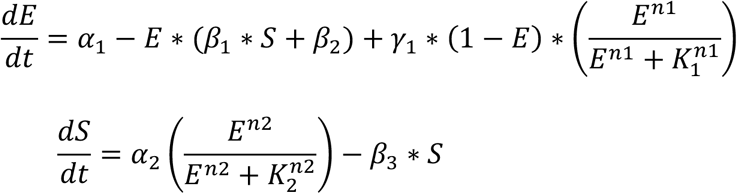

Here, *E* and *S* represent the activator and inhibitor respectively. Here, *β*_1_ is the inhibitor-controlled inactivation rate and *β*_2_ is the autonomous inactivation rate. *γ*_1_ controls the non-linear positive feedback for the activator which is represented as a Hill function. For the inhibitor equation, *α*_2_ is the activator dependent production rate and *β*_3_ is the inhibitor autonomous inactivation rate. We have assumed that the activator influences inhibitor production in a Hill function like manner as well.

We assume *α*_1_, the activator production rate, increases linearly in this system and solve the system of equations using ODE23s in MATLAB. The behavior of the nullclines as a function of *α*_1_ is plotted in Figure S2F with corresponding trajectories. The other parameters are set at *α*_2_ = 0.4, *β*_1_ = 3, *n*_1_ = 8, *n*_2_ = 8, *K*_1_ = 0.42, *K*_2_ = 0.47, *γ*_1_ = 2.5, *β*_2_ = 1.0, *β*_3_ = 0.3. In this model, with increase in *α*_1_, a stable node and saddle collide and then disappear generating oscillations. Subsequent increases in *α*_1_ is accompanied by a decrease in the time-period.

### Extraction of spry2 expression

Max projections from the transgenic line *TgKI(spry2-hsp70l:venus-Pest)* were difficult to analyze for notochord cells because of signal from the surrounding tissues (Figure S3B). To extract out the notochord, best match using cross-correlation were made between sections along the Y-Z axis and generated circles with varying radii and centers. The process was applied every 25 slices and the extracted parameters of the circle were separately fit to the X axis using least-squares. This then allowed the extraction of the extent of the notochord tube for each position on the X axis, any other signal beyond the fitted circles were removed, along with cutting the circle in half to remove part of the medial-lateral signal. Individual tiles were evaluated separately, then stitched together using cross-correlation over overlapping regions. Detected cells were then segmented with Cellpose.

### Quantifying spry2 and dusp6 knockdown experiments

F0 injected or non-injected controls were housed in similar conditions until 8 dpf. The two groups were first matched by length to only allow for comparisons between similarly developed larvae (Figure S6A). A region encompassing segments 6-8 (∼ 400 *μm*) was selected for further analysis through confocal microscopy. After extracting total *entpd5a:pKRed* fluorescence, a threshold near the 25^th^ percentile of non-injected fluorescence values was set to identify any injected larvae that would fall below and thus show a phenotype in the segmentation process. Along with the entire control group, these subset of injected larvae were then monitored for Erk activity at two different time points separated by roughly 5 hours. All activity per group was then pooled and the distributions were analyzed. Since the lower half of the distributions showed the most difference, values in the two groups below their respective medians were compared further and statistics were calculated using a Mann-Whitney U Test.

### In Situ Hybridization Chain Reaction (HCR)

Protocol for performing HCR was adopted from other published methods^54^. HCR Probes were obtained from IDT (Oligo Pools, oPools). Larvae at 3 and 5 dpf were anesthetized in 10X tricaine for 30 mins, then fixed in 4% PFA at 1X PBS for 2 hours at room temperature. Following two 1X PBS washes, the samples were then added to acetone for 20 mins and stored at −20 °C. After performing 3 PBS washes, the samples were pre-hybridized in probe hybridization buffer for 30 mins and then incubated in solution containing 0.5 pmol of each probe in 125*μL* at 37 °C for 40 hours. Following hybridization, larvae were washed in a probe wash buffer followed by two 5X SSCT washes for 10 minutes. The next step was addition to the amplification buffer, first only the samples were incubated for 30 mins. Then 7.5 pmol of hairpin 1 and 2 were separately snap cooled (heated to 95 °C for 90 seconds and cooled to room temperature) and added to 125*μL* of amplification buffer. Samples were then left overnight in a dark compartment. Hairpin solution was washed out the next day with 5X SSCT and mounted for confocal microscopy.

Data quantification was performed by first thresholding *egf* expression to eliminate background, and the resulting masks were then applied to the original image to recover true intensity values. Same threshold was applied for different developmental stages at 3 and 5 dpf across a technical replicate and similar regions towards the posterior of the larvae were used for quantification. Signal in the *col2* channel was then used to estimate the notochord, by building convex hulls across the different 3D planes. Intensity from *egf* expression was then summed up for each 3D slice in the estimated notochord region. To account for variability between experiments, all larvae from a technical replicate were normalized to their corresponding combined mean 3dpf and 5dpf *egf* measurement.

### Drug Treatments

#### Frequency Change

8 dpf larvae containing *cmn-hsp70l:erk-ktr-eGFP*;*entpd5a:pKRed* were pre-imaged. Larvae displaying high Erk activity were subsequently followed. They were immersed in 2 *μ*M PD0325901(Selleck Chem:S1036) or equivalent amount of DMSO (Sigma-Aldrich: D8418) in fish system water for about 2 hours. This was then washed out, by rinsing the larvae 3 times in system water. Following a post-treatment image, larvae were imaged every 4 hours while monitoring signs of high Erk activity, indicating the next occurrence of an oscillation cycle. If any high activity was detected, another image was taken within the next 30 mins to capture the peak of the oscillation cycle.

#### Continuous Treatment

3 dpf transgenic larvae containing *col9a2:erk-ktr-mCerulean* were immersed in 50 *μ*M Afatinib/BIBW2992 (Selleck Chem: S1011) or an equivalent amount of DMSO in fish egg water for about 50 hours. Post treatment images were taken to assess Erk activity.

Larvae at 5dpf and 7 dpf expressing *entpd5a:pKRed* were immersed in 2 *μ*M PD0325901/ Mirdametinib (Figures 4D-4F) or 20 *μ*M Afatinib/BIBW2992 (Figures S9E-S9F) for about 50 hours. An equivalent amount of DMSO in fish system water was used for each control group. Drug solutions were replaced about every 24 hours. To assess changes in individual segment fluorescence, different segment regions were tracked between the pre and post images (Methods: *entpd5a* fluorescence extraction). Any cell clusters in the post-treated image which could not be assigned to a previous position in the pre-treated image were marked as new cells added to segments. To account for differences in imaging time within an experiment, the number of cells added or changes in fluorescence per segment, were rescaled by the exact time differences for each larva between pre and post treated images.

### Pulsed treatments

Larvae expressing *entpd5a:pKRed* at 7 dpf evening or 8 dpf morning were immersed in 2 *μ*M PD0325901/ Mirdametinib for a 10-hour duration. This was done twice with an additional 10 hour recovery period in between, in which the larvae were kept in fish water. Total fluorescence of *entpd5a:pKRed* per larva was monitored roughly 20 hours apart, before the onset of each pulse, along with post treatment images taken after the drug or DMSO solutions were rinsed out following the second pulse. They were imaged one final time, about 10 hours later. Only larvae displaying similar initial levels of total *entpd5a:pKRed* fluorescence were chosen for further analysis (Figure S9G)

### Statistical Analysis

All statistical tests were performed in Python (v3.10.7) using scipy (v1.11.1) and statsmodels (v0.15.0). Two independent groups were compared using a two-sided Mann-Whitney U test (Figures 3, 4, S6, S9) while nested cellular measurements within larvae (Figures S3 and S5) were compared using a linear mixed-effects model. Significance values were noted as follows: \**p* < 0.05, ** *p <* 0.01, *** *p* < 0.001 and ns (not significant). Exact p-values reported in each legend.

## SUPPLEMENTARY FIGURES

**Figure S1.**
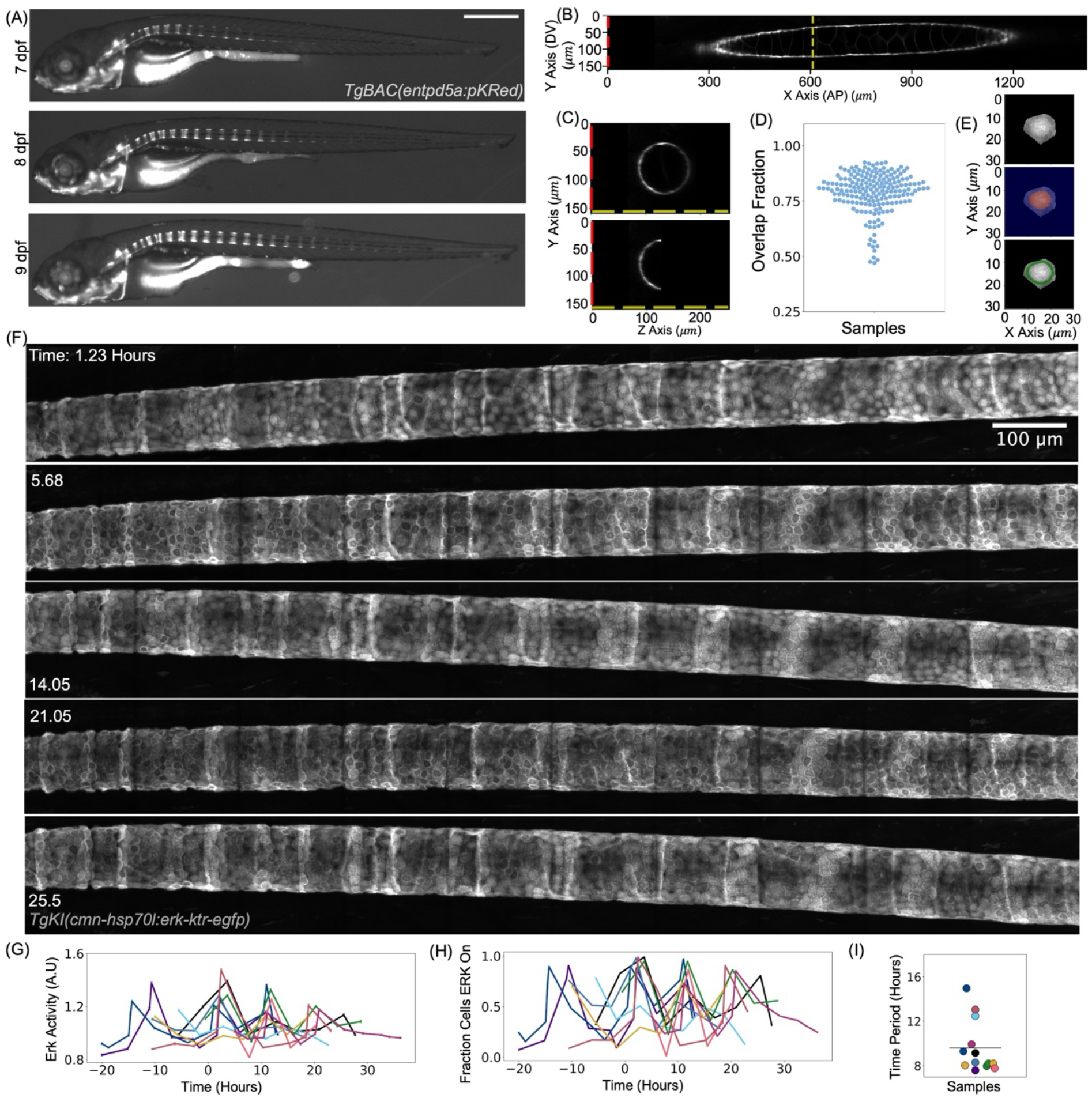
Characterization of segment progression and Erk sensor activity. Related to Figure 1. (A) Raw data of an entire larva expressing *entpd5a:pKRed* imaged between 7-9 dpf. New segments are added gradually to the notochord. Scale bar 500 *μm*. (B) Example section of the notochord as visualized along the Dorsal-Ventral (Y) and Anterior Posterior (X) axes at a particular position on the medial-lateral (Z) axis. (C) Top, cross sections in the Y-Z plane at a particular X position. Bottom, cross section after removing half of the signal along the Z axis. (D) Intersection over Union metric quantified in masks obtained from a trained model in Cellpose compared to a test set of manually identified masks. (E) Example cell with estimated nuclei pseudo-colored in red, followed by marking an outer ring of 3 pixels where the cytoplasmic intensity was evaluated. (F) Raw data of Erk sensor activity reported by *cmn-hsp70l:erk-ktr-eGFP* used to quantify Figure 1G. Scale bar 100 *μm*. (G) Combining n = 10 oscillator timecourses (N = 7 independent experiments) taken at 7-8 dpf. Data is represented as averaged Erk Activity± SEM and timecourses have been aligned for visualization. (H) Fraction of cells with high Erk activity plotted over time. This fraction is calculated by counting all cells whose cytoplasmic to nuclear intensity is greater than 1 compared to the whole population. (I) Time period of oscillations calculated by measuring trough-trough distances from individual larval time-courses in (H) (number of cycles = 13, horizontal line indicates mean).

**Figure S2.**
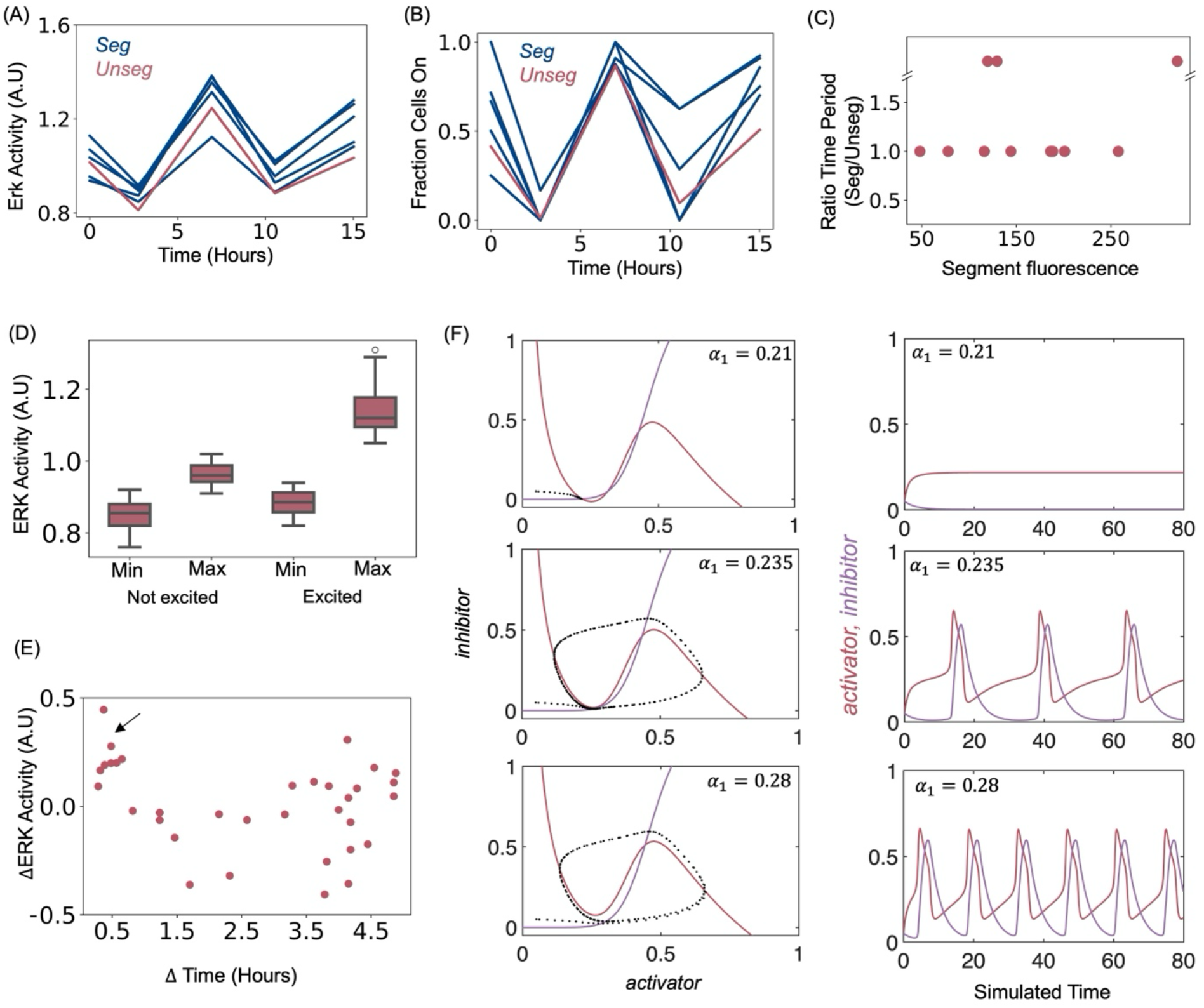
Characterizing Erk activity through a SNIC bifurcation. Related to Figure 2. (A) Erk activity as a function of time, separated in segmented and unsegmented regions, demonstrates that these regions have similar behaviors (n = 1, number of segments = 5). (B) Fraction of cells with high Erk activity quantified in different segmented and unsegmented regions. Same larva as in (A). (C) Ratio of time periods between segmented and unsegmented domains plotted as a function of total segment fluorescence (n = 2 larvae from N = 2 independent experiments, number of segments = 12); points above the break indicate regions where time-period measurement couldn’t be correctly estimated within our sampling window. (D) Minimum and maximum Erk activity seen in cell trajectories from Figure 2F before the bifurcation point. A cell is categorized as excitable if it crosses a threshold of high Erk activity (1.02) before coming down. (E) Change in Erk activity plotted against time interval between subsequent imaging time-points; spikes in Erk activity are seen near the oscillation peak for more frequently sampled points (black arrow). Data is quantified in n = 4 larvae from N = 3 independent experiments. (F) Left Column: Nullclines of the activator-inhibitor system as a function of activator production rate *α*_1_. For low *α*_1_, the nullclines intersect at three positions on the phase plane, creating one stable and two unstable fixed points. As *α*_1_ is increased, the stable and leftmost unstable fixed points collide. Subsequent increase in *α*_1_ leads to disappearance of this steady state, generating oscillations. Right Column: Corresponding time-series of the activator-inhibitor variable as a function of *α*_1_. Oscillations become faster with increasing *α*_1_. Other simulation parameters listed under Methods.

**Figure S3.**
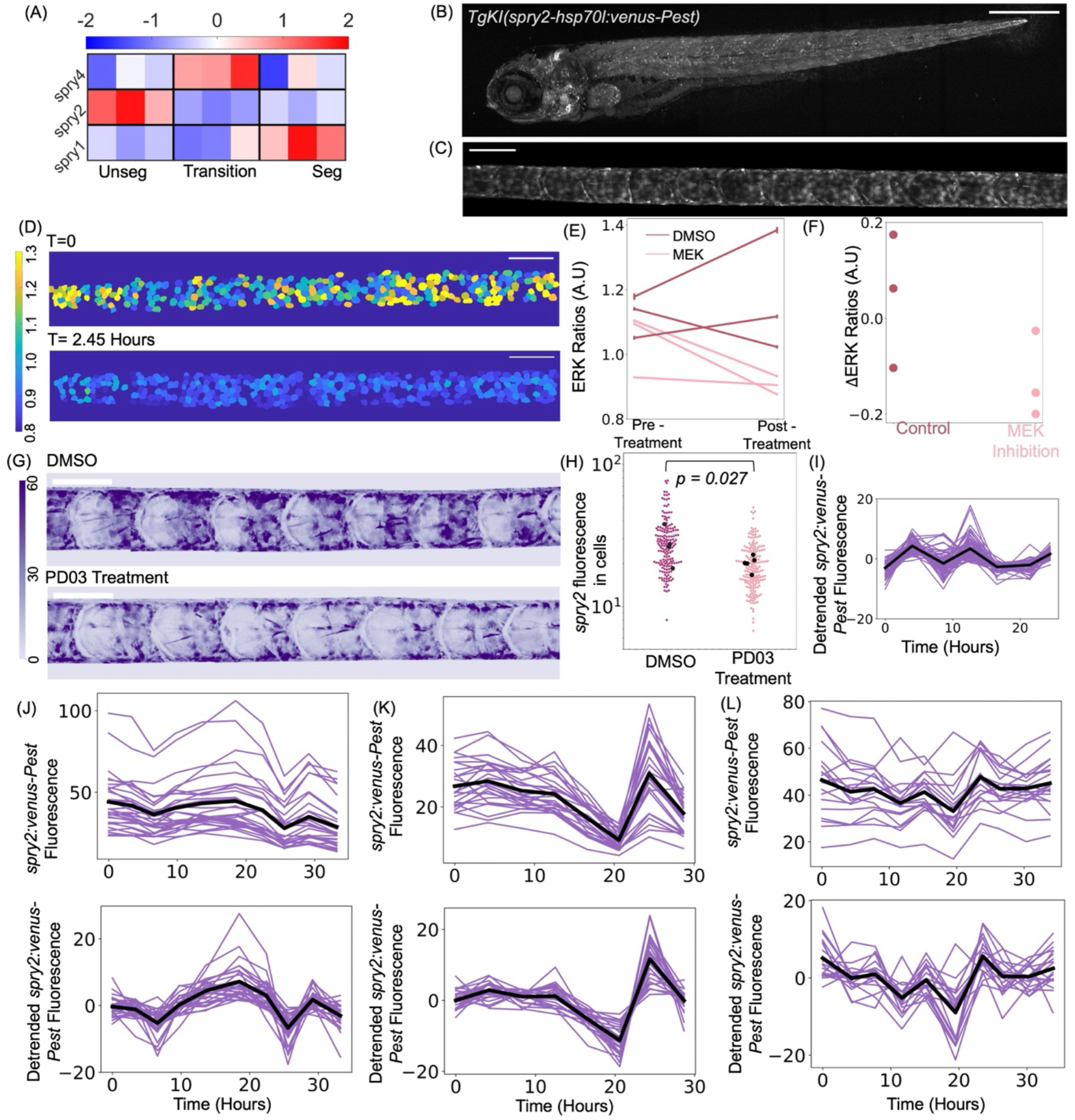
Visualizing *spry2* expression in the zebrafish notochord. Related to Figure 3. (A) z-score calculated for *sprouty* genes from a transcriptomic dataset collected at 13 dpf. (B) Image showing an entire 5 dpf larva expressing *TgKI*(*spry2-hsp70l:venus-Pest)* (Scale bar 500 *μm*). (C) Computational extraction of the notochord region in a 6dpf *TgKI*(*spry2-hsp70l:venus-Pest)* larva. Scale bar 100 *μm*. (D) Heat maps showing reduction of Erk activity in Mek inhibitor treated (2 *μM*) larva within a 2-hour treatment window. Scale bar 100 *μm*. (E) Average Erk activity± SEM quantified pre and post treatment in control or treated larvae (n = 3 per condition quantified from one representative experiment). (F) Change in Erk activity quantified between the pre and post treatment images in (E). Average activity in treated goes down consistently, while the control group could show increase or decreases depending on the pre-treatment phase of the oscillation cycle. (G) Fluorescence intensity quantified as a color map following a Mek inhibition in 8 dpf larvae expressing *spry2-hsp70l:venus-Pest*. Scale bar 100 *μm*. (H) *spry2-hsp70l:venus-Pest* expression quantified in individual cells. Black dots represent larval averages; N = 1, n = 5 treated larvae with 187 cells and n = 4 control larvae with 174 cells. Comparisons made on log transformed fluorescence values using a linear-mixed model with treatment as the fixed effect, *p = 0.027.* (I) Detrended *spry2-hsp70l:venus-Pest* fluorescence for the larva as shown in Figure 3B. (J-L) Each column represents a different larva expressing *spry2-hsp70l:venus-Pest.* Top row shows total fluorescence levels and bottom shows detrended levels. n = 29 cells in (J), n = 27 cells in (K) and n = 19 cells in (L). Black lines represent the average of all the traces.

**Figure S4.**
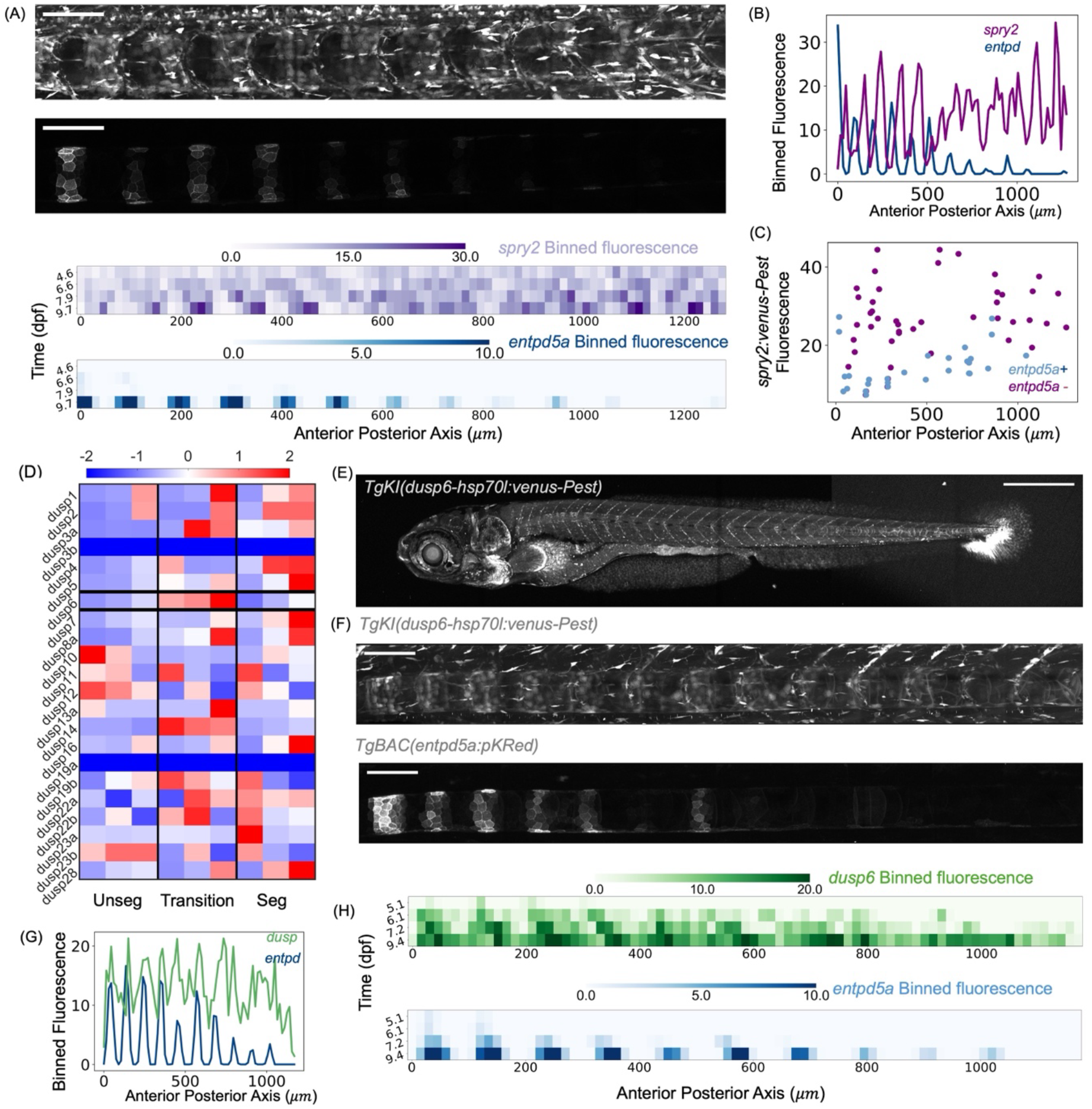
Segmented patterns of negative feedback pathway components. Related to Figure 3. (A) Top: Example larva at 8 dpf expressing *spry2-hsp70l:venus-Pest* and *entpd5a:pKRed*. Scale bar 100 *μm*. Bottom: Binned *spry2* and *entpd5a* fluorescence levels in 15 *μm* bins in a larva followed between 4-9 dpf, with colorbars representing the intensities. (B) Binned profile trace of *spry2* fluorescence (purple) at 9 dpf across the Anterior-Posterior axis. (C) Time averaged *spry2:venus-Pest* expression as a function of position along the AP axis quantified in *entpd5a+* (blue dots) and *entpd5a-*(purple dots) populations. (D) z-score calculated for *dusp* genes from a transcriptomic dataset (same dataset as shown in Figure S3A) collected at 13 dpf. (E) Image showing an entire 6 dpf larva expressing *TgKI*(*dusp6-hsp70l:venus-Pest)* (Scale bar 500 *μm*). (F) Image showing a 7 dpf larva expressing *TgKI(dusp6-hsp70l:venus-Pest*) along with *entpd5a:pKRed* (Scale bar 100 *μm*). (G) Binned profile trace of *dusp6* fluorescence at 9 dpf (green) across the Anterior-Posterior axis. (H) Binned *dusp6* and *entpd5a* fluorescence levels in 15 *μm* bins in a larva followed between 5-9 dpf, with colorbars representing the intensities.

**Figure S5.**
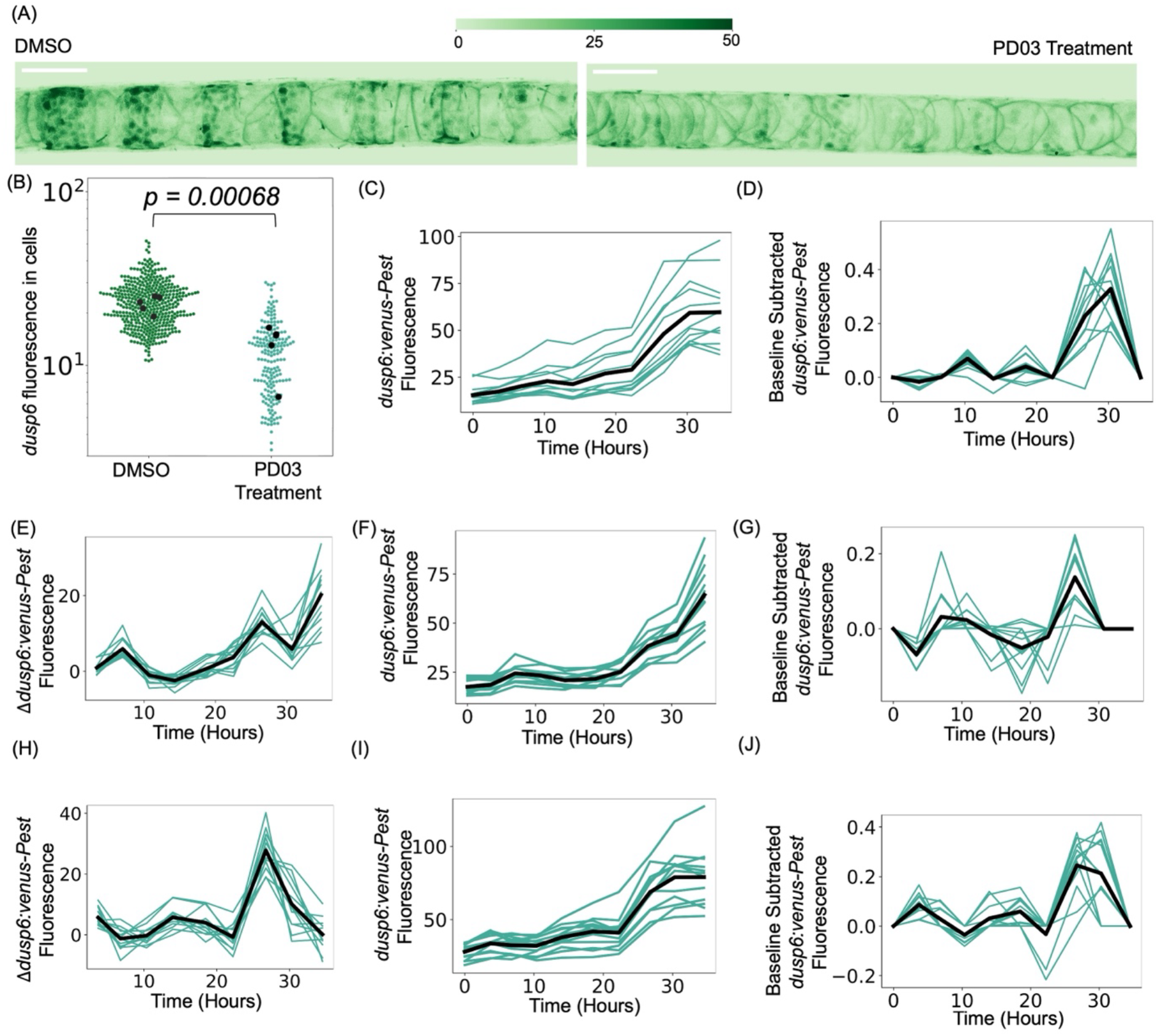
Visualizing *dusp6* expression over time in the zebrafish notochord. Related to Figure 3. (A) Fluorescence intensity quantified as a color map following a Mek inhibition in 8 dpf larvae expressing *dusp6-hsp70l:venus-Pest*. Scale bar 100 *μm*. (B) *dusp6-hsp70l:venus-Pest* expression quantified in individual cells following treatment: N=1, DMSO: n = 5 larvae, 418 cells; PD03: n = 5 larvae, 190 representing all cells that could be detected post treatment. Comparisons were performed using a linear mixed-effects model with treatment as the fixed effect, *p* = 0.00068. (C-D) Total and baseline subtracted fluorescence levels in a larva expressing *dusp6-hsp70l:venus-Pest* over time. Same sample as shown in Figure 3E. (E-J) Each row is a different larva showing timepoint-by-timepoint differences in *dusp6* expression, total and baseline subtracted fluorescence levels. Number of cells = 12 for each larva.

**Figure S6.**
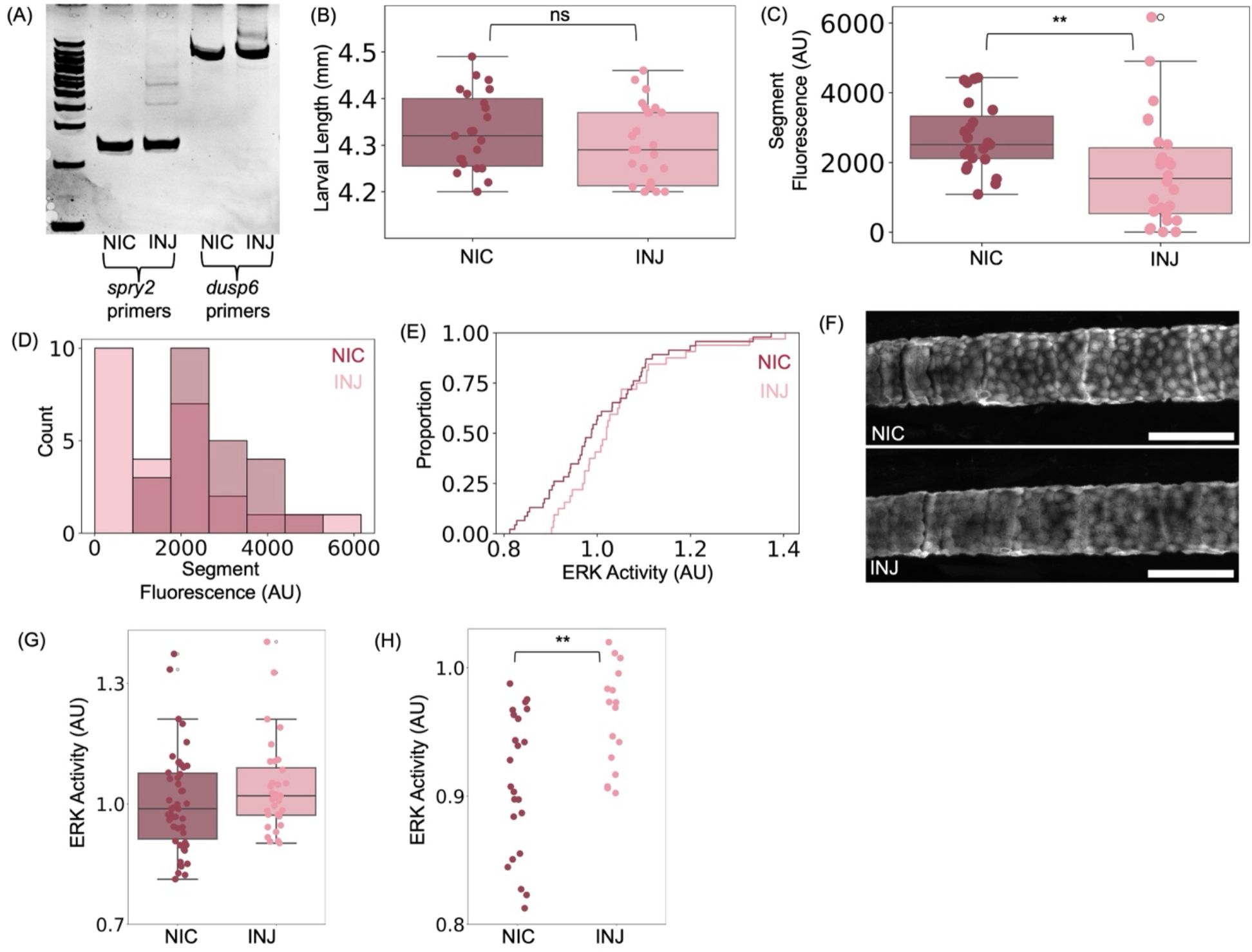
sgRNA pools targeting *spry2* and *dusp6.* Related to Figure 3. (A) PCR analysis showing gRNA efficiency. (B) Length distributions of Non-Injected Controls (NIC) and Injected 8dpf larvae; N = 3 independent experiments; n = 26 injected larvae, n = 23 larvae for NIC; Mann-Whitney U test, *p* = 0.29 (ns). Individual dots represent individual larvae. (C) Total segment fluorescence quantified from *entpd5a:pKRed* expression in controls and injected larvae; Mann-Whitney U test, *p* = 0.0043. Imaged region covers segments 6-8. (D) Distribution from (C) quantified as histograms. (E) Empirical Cumulative Distribution Function (ECDF) of Erk activity in larvae displaying segmentation phenotype (n = 16), selected by setting a threshold of total *entpd5a* fluorescence lower than 2000 A.U compared to n = 23 control larvae. Erk is imaged at two separate points for each larva, roughly 5 hours apart and all data is pooled together per condition. (F) Example of lowest Erk activity attained amongst either the non-injected controls or injected larvae. Represents lowest rank from the ECDF plot. Scale bar 100 *μm*. (G) Distribution of Erk activity in the two conditions (n injected = 16, n controls = 23). (H) Comparison of Erk activity levels below each condition’s own median for NIC and injected larvae. Data shown as a scatter plot were derived from measurements in (G), yielding n = 16 injected and n = 23 control data points; Mann-Whitney U test, *p* = 0.0045.

**Figure S7.**
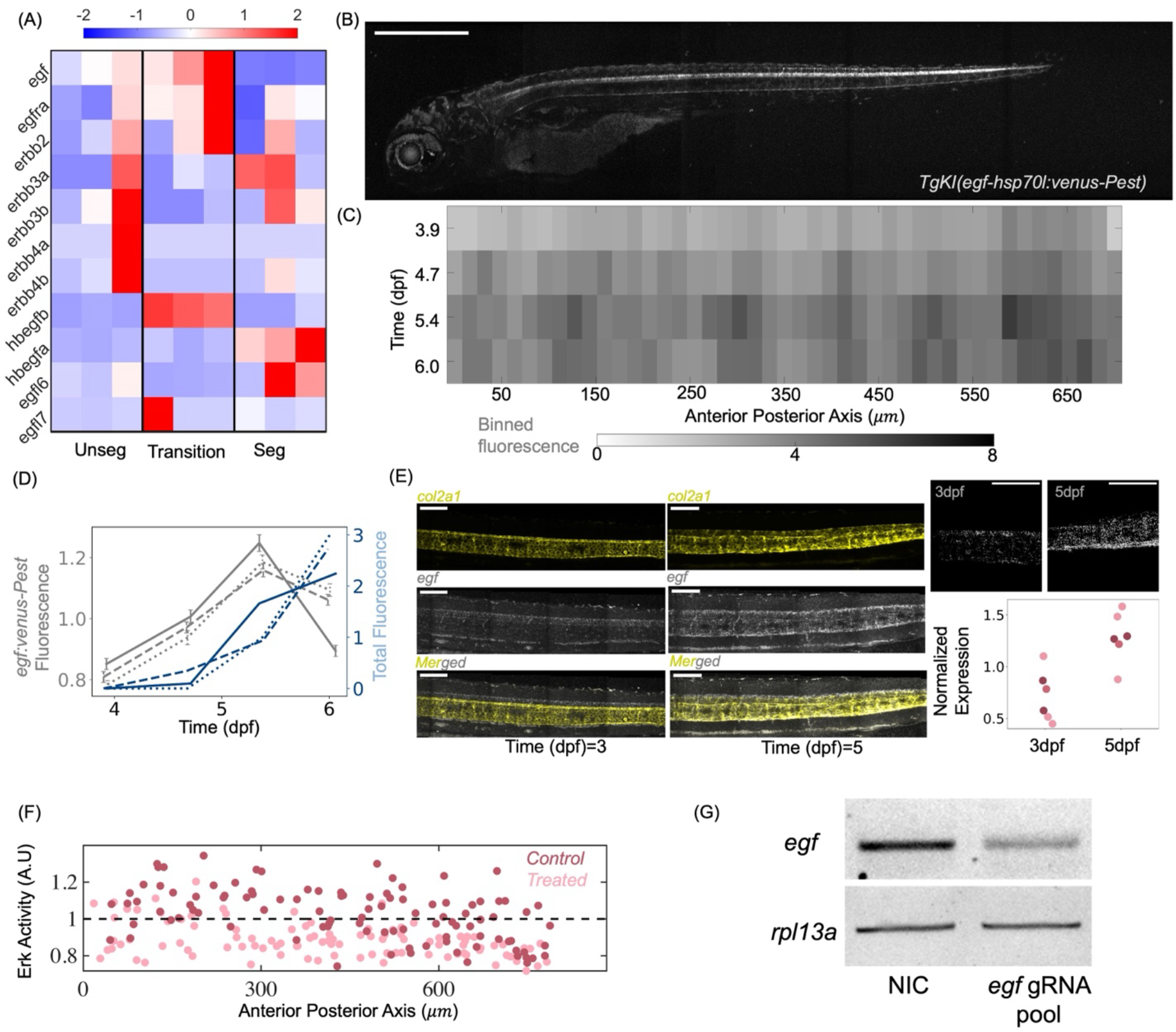
Dynamics of *egf* expression in the notochord. Related to Figure 3. (A) z-scores calculated for genes associated with the *egf* pathway from a transcriptomic dataset (same dataset as shown in Figures S3A and S4D) collected at 13 dpf. (B) Image showing an entire 5 dpf larva expressing *TgKI*(*egf-hsp70l:venus-Pest*) (Scale bar 500 μm). (C) Binned expression of the *egf* reporter in 15 *μm* bins monitored through time. Colorbar represents intensity levels. (D) Average ± SEM *egf-hsp70l:venus-Pest* and *entpd5a*:pKRed fluorescence levels as a function of time. This analysis indicates an increase in *egf* expression around 5 dpf, similar to when the first bursts of *entpd5a* expression can be detected. (E) HCR in-situ comparing *egf* expression in 3 and 5 dpf larvae with *col2a1* expression for marking the notochord (Scale bar 100 μm). N = 3 independent experiments with n = 6 larvae total at each 3 and 5 dpf timepoint (n = 3, n = 2, n = 1 per experiment). Expression levels normalized within each experiment to the combined mean expression levels from both 3 and 5 dpf for visualization. Fold change calculated as the ratio between 5dpf to 3dpf levels are 1.91, 1.78 and 1.55 for each experiment. (F) Erk levels in individual cells as function of position along the AP axis following an EGFR inhibition by treating 3 dpf larvae for 50 hours with Afatinib at 50 *μM* (n = 3 larvae per condition shown from one experiment from Figure 3H). (G) RT-PCR to test gRNA efficiency.

**Figure S8.**
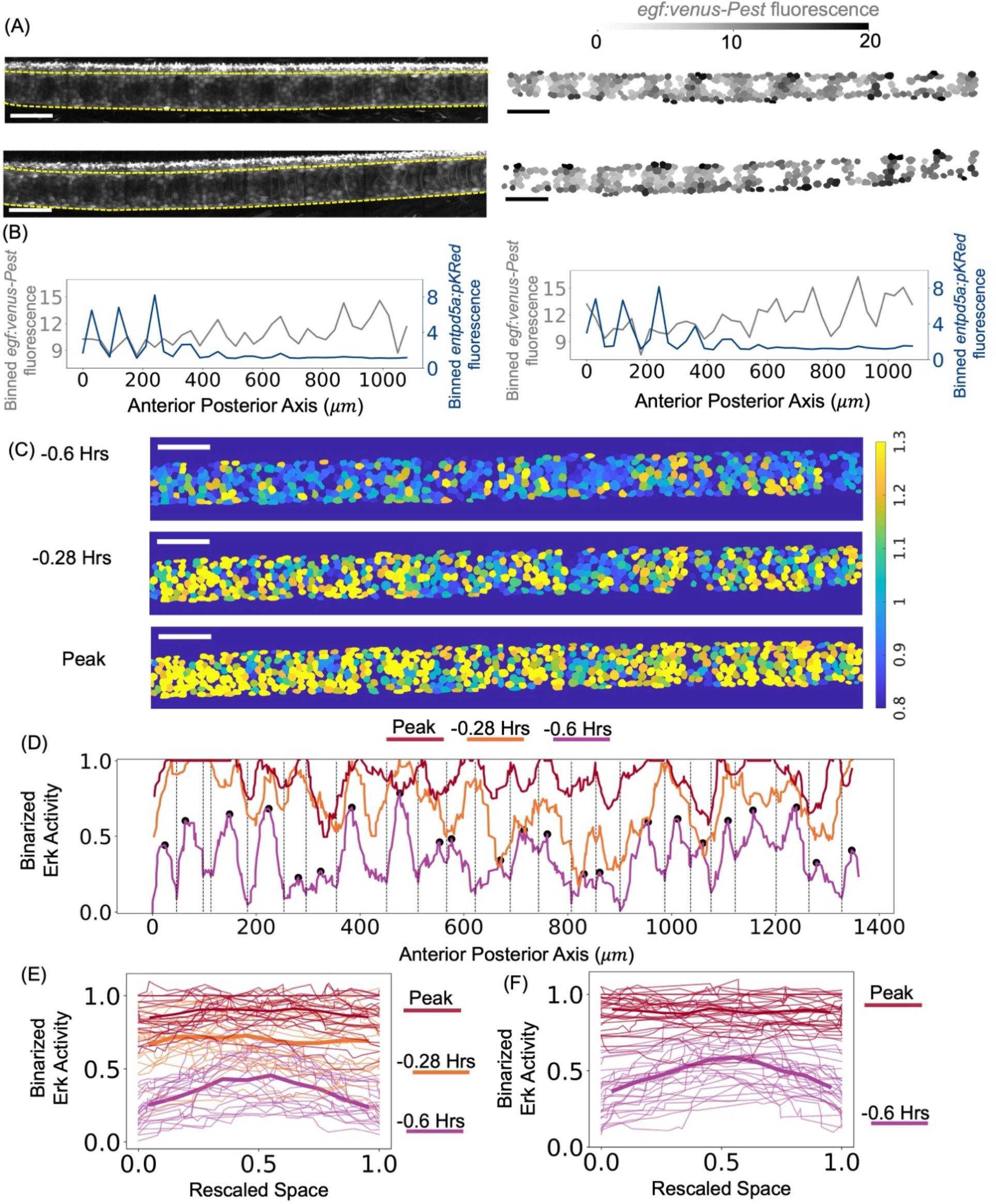
Segmented patterns of *egf* expression. Related to Figure 3. (A) 8dpf larvae expressing *TgKI*(*egf-hsp70l:venus-Pest*) on the left with segmented *venus-Pest*+ cells on the right with fluorescence levels mapped by the colorbar (n = 2 larvae shown from one experiment). Scale bars 100 μm. (B) Binned profile traces of the segmented images across the Anterior-Posterior axis with binned expression from *entpd5a*:pKRed in the same larva for comparison. (C) Heatmaps of Erk activity at time-points close to an oscillation peak. Time labels indicate the duration prior to the peak. Scale bar 100 μm. (D) Erk activity along the AP axis at time-points close to an oscillation peak quantified in n = 1 larva with legend for colors above. Activity for each cell is first scored as either 1 or 0 depending on whether Erk sensor cytoplasmic fluorescence is greater or less than nuclear fluorescence respectively. Then the profiles are passed through a Savitzky-Golay filter to indicate any local regions with increased activity (black dots). (E) Plot shows that aligning regions in (D) over time (boundaries indicated by dotted black lines), flatten out spatial variations close to the peak. (F) Additional example from different larva characterizing spatial variations close to an oscillation peak.

**Figure S9.**
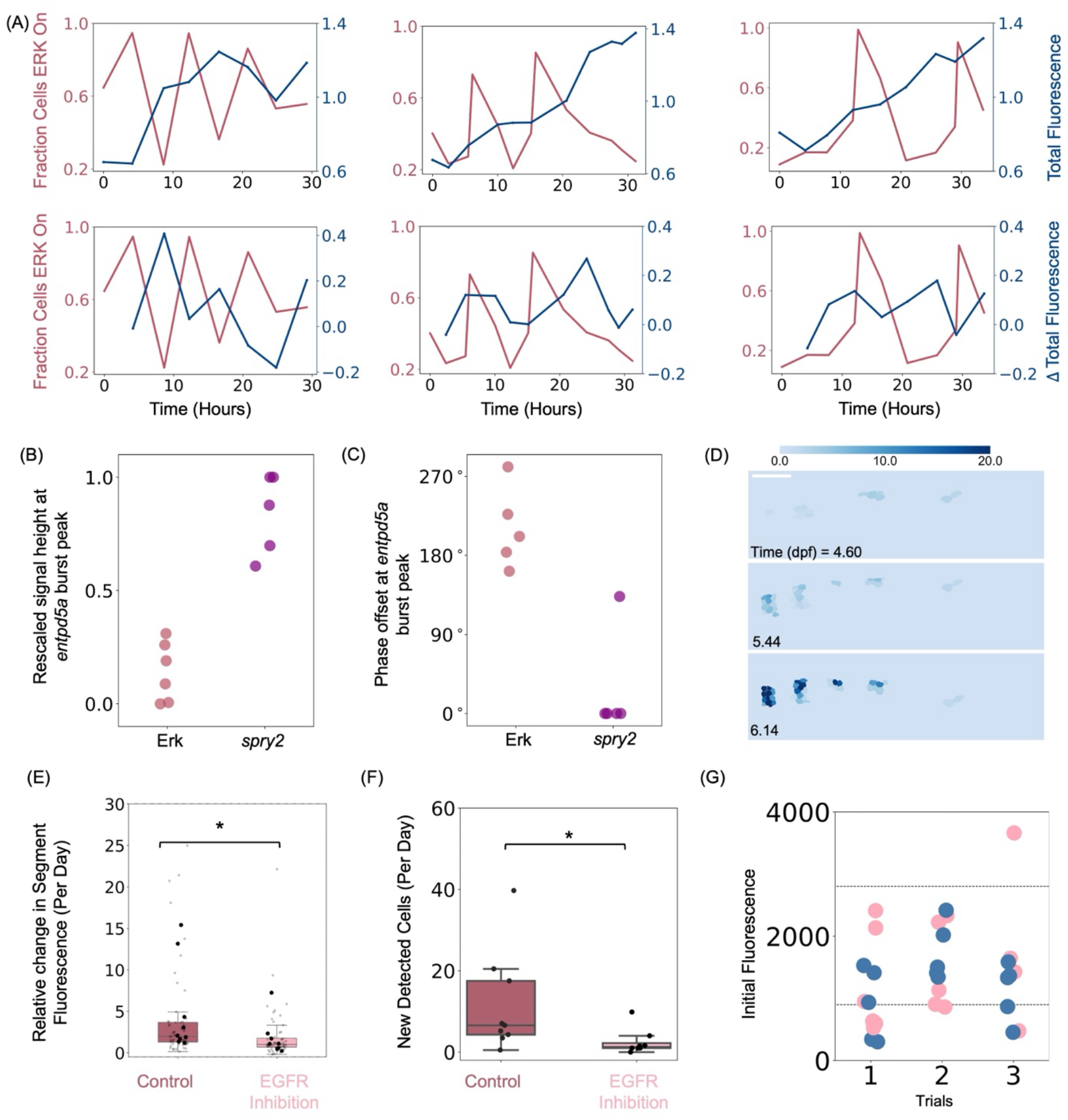
Inhibition of Erk activity. Related to Figures 4-5. (A) Additional examples with Erk sensor and *entpd5a* in the same larva (n = 3 larvae from N = 2 independent experiments). Top: Fraction of cells with high Erk activity and total *entpd5a:pKRed* fluorescence normalized by mean levels as a function of time in the same larva. Bottom: Fraction of cells with high Erk activity and timepoint-by-timepoint difference in *entpd5a:pKRed* fluorescence as a function of time in a single larva. (B) Estimate of signal height at the time-point a burst in *entpd5a:pKRed* expression is detected. Data for Erk activity timecourses is from Figure 4A and panels A, data for timecourses expressing *spry2-hsp70l:venus-Pest* from Figure 3C. Signal height is rescaled between the minimum and maximum values for each respective time-course. (C) Phase offset of a *entpd5a:pKRed* burst between encompassing peaks of Erk activity or *spry2-hsp70l:venus-Pest* expression. Offsets are calculated by measuring fractional distances from the leftmost signal peak to the *entpd5a* burst compared to the distance between the two signal peaks. Any offsets of 360° are converted back to 0°. Only *entpd5a* bursts detected above a threshold (0.15) are analyzed further. (D) Heatmap quantification of *entpd5a:pKRed* around 5 dpf shows the first forming segments. Scale bar 100 μm. (E) Relative change in fluorescence of existing segments following EGF receptor signaling inhibition pharmacologically at 7dpf. Small points indicate segment-level measurements, large points indicate per-larva means on which statistical comparison was performed (Mann-Whitney U test, *p =* 0.046). (F) Number of newly added cells in control and treated larvae from (E) (Mann-Whitney U test, *p* = 0.027). Larvae in (E-F) were treated to 20 *μM* Afatinib over 50 hours; N = 3 independent experiments, n = 9 control larvae, n = 8 treated larvae. (G) Total fluorescence of *entpd5a:pKRed* at pre-treatment for different experimental trials used in Figure 5F. Dashed lines indicate the fluorescence intervals used for selecting larvae for further analysis.

## Supplementary Information

**Video S1** Heat Maps of Erk Activity Show Oscillation Cycles Top: Oscillations in an 8dpf larva plotted as a heat map over 30 hours. Related to Figure 1G (see associated colorbar). Scale bar 100 μm. Bottom: Corresponding raw data of Erk sensor activity. Related to Figure S1F. Scale bar 100 μm.

**Video S2** Heat Maps Showing Increase in Erk Activity around 5 dpf Heat maps of Erk activity between 3-5dpf. Related to Figure 2E (see colorbar). Scale bar 100 μm.

**Video S3 Heat Maps of Erk Activity in a Control Larvae Following a Washout Experiment** Images of Erk activity following an 8dpf larva post DMSO treatment. Related to Figure 5A (see associated colorbar). Scale bar 100 μm.

**Video S4** Heat Maps of Erk Activity in a Mek Inhibited Larvae Following a Washout Experiment Images of Erk activity following an 8dpf larva post Mek inhibition. Related to Figure 5A (see associated colorbar). Scale bar 100 μm.

## References

1. Palmeirim I., Henrique D., Ish-Horowicz D. & Pourquié, O. Avian hairy gene expression identifies a molecular clock linked to vertebrate segmentation and somitogenesis. Cell 91, 639–648 (1997).

2. Hubaud, A. & Pourquie, O. Signalling dynamics in vertebrate segmentation. Nat. Rev. Mol. Cell. Biol. 15, 709–721 (2014).

3. Cooke, J. & Zeeman, E.C. A clock and wavefront model for control of the number of repeated structures during animal morphogenesis. J. Theor. Biol. 58, 455–476 (1976).

4. Negrete, J., Jr. & Oates, A. C. Towards a physical understanding of developmental patterning. Nat. Rev. Genet. 22, 518–531 (2021).

5. Fleming, A., Kishida, M. G., Kimmel, C. B. & Keynes, R. J. Building the backbone: the development and evolution of vertebral patterning. Development 142, 1733–1744 (2015).

6. Fleming, A., Keynes, R. & Tannahill, D. A central role for the notochord in vertebral patterning. Development 131, 873–880 (2004).

7. Wopat, S. et al. Spine Patterning Is Guided by Segmentation of the Notochord Sheath. Cell. Rep. 22, 2026–2038 (2018).

8. Lleras Forero, L., et al. Segmentation of the zebrafish axial skeleton relies on notochord sheath cells and not on the segmentation clock. eLife 7 (2018).

9. Pogoda, H. M., et al. Direct activation of chordoblasts by retinoic acid is required for segmented centra mineralization during zebrafish spine development. Development 145 (2018).

10. Wopat, S., et al. Notochord segmentation in zebrafish controlled by iterative mechanical signaling. Dev. Cell. 59, 1860–1875 (2024).

11. Peskin, B., et al. Dynamic BMP signaling mediates notochord segmentation in zebrafish. Curr. Biol. 33, 2574–2581 (2023).

12. Pogoda, H. M., Riedl-Quinkertz, I. & Hammerschmidt, M. Direct BMP signaling to chordoblasts is required for the initiation of segmented notochord sheath mineralization in zebrafish vertebral column development. Front. Endocrinol. (Lausanne) 14, 1107339 (2023).

13. Holley S.A., Geisler R. & Nüsslein-Volhard, C. Control of her1 expression during zebrafish somitogenesis by a delta-dependent oscillator and an independent wave-front activity. Genes. Dev. 14, 1678–1690 (2000).

14. Sarrazin A.F., Peel A.D. & Averof, M. A segmentation clock with two-segment periodicity in insects. Science 336, 338–341 (2012).

15. El-Sherif, E., Averof, M. & Brown, S. J. A segmentation clock operating in blastoderm and germband stages of Tribolium development. Development 139, 4341–4346 (2012).

16. Pascoal, S., et al. A molecular clock operates during chick autopod proximal-distal outgrowth. J. Mol. Biol. 368, 303–309 (2007).

17. Chou, K. T., et al. A segmentation clock patterns cellular differentiation in a bacterial biofilm. Cell 185, 145–157 (2022).

18. Richmond, D. L. & Oates, A. C. The segmentation clock: inherited trait or universal design principle? Curr. Opin. Genet. Dev. 22, 600–606 (2012).

19. Dequéant, M.L., et al. A complex oscillating network of signaling genes underlies the mouse segmentation clock. Science 314,1595–1598 (2006).

20. Niwa, Y., et al. Different types of oscillations in Notch and Fgf signaling regulate the spatiotemporal periodicity of somitogenesis. Genes. Dev. 25, 1115–1120 (2011).

21. Simsek, M. F., et al. Periodic inhibition of Erk activity drives sequential somite segmentation. Nature 613, 153–159 (2023).

22. De Simone, A., et al. Control of osteoblast regeneration by a train of Erk activity waves. Nature 590, 129–133 (2021).

23. Regot, S., Hughey, J. J., Bajar, B. T., Carrasco, S. & Covert, M. W. High-sensitivity measurements of multiple kinase activities in live single cells. Cell 157, 1724–1734 (2014).

24. Peskin, B., et al. Notochordal Signals Establish Phylogenetic Identity of the Teleost Spine. Curr. Biol. 30, 2805–2814 (2020).

25. Garcia, J., et al. Sheath Cell Invasion and Trans-differentiation Repair Mechanical Damage Caused by Loss of Caveolae in the Zebrafish Notochord. Curr. Biol. 27, 1982–1989 (2017).

26. Pikovsky, A., Rosenblum, M., & Kurths, J. Synchronization: A Universal Concept in Nonlinear Sciences. Cambridge University Press (2001).

27. Strogatz, S.H. From Kuramoto to Crawford: Exploring the Onset of Synchronization in Populations of Coupled Oscillators. Physica D 143, 1–20 (2000).

28. Strogatz, S.H. Nonlinear Dynamics and Chaos: With Applications to Physics, Biology, Chemistry, and Engineering. CRC Press, ed.3 (2024).

29. Izhikevich, E.M. Dynamical Systems in Neuroscience: The Geometry of Excitability and Bursting. The MIT Press (2010).

30. Murayama, Y., et al. Low temperature nullifies the circadian clock in cyanobacteria through Hopf bifurcation. Proc. Natl. Acad. Sci. USA 114, 5641–5646 (2017).

31. Salvi, J. D., D, O. M. & Hudspeth, A. J. Identification of Bifurcations from Observations of Noisy Biological Oscillators. Biophys. J. 111, 798–812 (2016).

32. Sanchez, P. G. L., et al. Arnold tongue entrainment reveals dynamical principles of the embryonic segmentation clock. eLife 11 (2022).

33. Tsai, T. Y., et al. Robust, tunable biological oscillations from interlinked positive and negative feedback loops. Science 321, 126–129 (2008).

34. Tyson, J.J., Chen, K.C. & Novak, B. Sniffers, buzzers, toggles and blinkers: dynamics of regulatory and signaling pathways in the cell. Curr. Opin. Cell. Biol. 15, 221–231 (2003).

35. Ferrell, J. E., Jr., Tsai, T. Y. & Yang, Q. Modeling the cell cycle: why do certain circuits oscillate? Cell 144, 874–885 (2011).

36. Hanafusa, H., Torii, S., Yasunaga, T. & Nishida, E. Sprouty1 and Sprouty2 provide a control mechanism for the Ras/MAPK signalling pathway. Nat. Cell. Biol. 4, 850–858 (2002).

37. Li, C., Scott, D. A., Hatch, E., Tian, X. & Mansour, S. L. Dusp6 (Mkp3) is a negative feedback regulator of FGF-stimulated ERK signaling during mouse development. Development 134, 167–176 (2007).

38. Simsek, M. F., et al. The vertebrate segmentation clock drives segmentation by stabilizing Dusp phosphatases in zebrafish. Dev. Cell. 60, 669–678 (2025).

39. Jiang, YJ., Aerne, B., Smithers, L. et al. Notch signalling and the synchronization of the somite segmentation clock. Nature 408, 475–479 (2000).

40. Jutras-Dubé, L., El-Sherif, E., & François, P. “Geometric models for robust encoding of dynamical information into embryonic patterns,” eLife, 9 (2020).

41. Riedel-Kruse, I. H., Müller, C., & Oates, A. C. Synchrony dynamics during initiation, failure, and rescue of the segmentation clock. Science 317, 1911–1915 (2007).

42. Yoshioka-Kobayashi, K. et al. Coupling delay controls synchronized oscillation in the segmentation clock. Nature 580, 119–123 (2020).

43. Keynes, R., & Lumsden, A. Segmentation and the Origin of Regional Diversity in the Vertebrate Central Nervous System, Neuron 2, 1–9 (1990).

44. Wingert, R.A. et al. The Cdx Genes and Retinoic Acid Control the Positioning and Segmentation of the Zebrafish Pronephros. PLoS Genetics 3, 1922–1938 (2007).

45. Westerfield, M. The Zebrafish Book: A Guide for the Laboratory Use of Zebrafish (Danio rerio). University of Oregon Press, ed.5 (2007).

46. Parichy, D. M., Elizondo, M. R., Mills, M. G., Gordon, T. N. & Engeszer, R. E. Normal table of postembryonic zebrafish development: staging by externally visible anatomy of the living fish. Dev. Dyn. 238, 2975–3015 (2009).

47. Levic, D. S., Yamaguchi, N., Wang, S., Knaut, H., & Bagnat, M. Knock-in tagging in zebrafish facilitated by insertion into non-coding regions. Development 148 (2021).

48. Marchetti, S., et al. Extracellular Signal-Regulated Kinases Phosphorylate Mitogen-Activated Protein Kinase Phosphatase 3/DUSP6 at Serines 159 and 197, Two Sites Critical for Its Proteasomal Degradation. Mol Cell Biol, 25, 854–864 (2005).

49. Nichols, A., et al. Substrate recognition domains within extracellular signal-regulated kinase mediate binding and catalytic activation of mitogen-activated protein kinase phosphatase-3. Journal of Biological Chemistry, 275, 24613–24621 (2000).

50. Stringer, C., Wang, T., Michaelos, M. & Pachitariu, M. Cellpose: a generalist algorithm for cellular segmentation. Nat. Methods. 18, 100–106 (2021).

51. Cox, B. Investigating Dynamics of Tissue Regeneration via Live Imaging of Zebrafish Scales. Duke University (2019).

52. Pomerening, J. R., Kim, S. Y. & Ferrell, J. E., Jr. Systems-level dissection of the cell-cycle oscillator: bypassing positive feedback produces damped oscillations. Cell 122, 565–578 (2005).

53. Novák, B. & Tyson, J. J. Design principles of biochemical oscillators. Nat. Rev. Mol. Cell. Biol. 9, 981–991 (2008).

54. Munjal, A., Hannezo, E., Tsai, T. Y., Mitchison, T. J. & Megason, S. G. Extracellular hyaluronate pressure shaped by cellular tethers drives tissue morphogenesis. Cell 184, 6313–6325 (2021).

